# Tensin3 interaction with talin drives formation of fibronectin-associated fibrillar adhesions

**DOI:** 10.1101/2021.07.16.452612

**Authors:** Paul Atherton, Rafaella Konstantinou, Suat Peng Neo, Emily Wang, Eleonora Balloi, Marina Ptushkina, Hayley Bennett, Kath Clark, Jayantha Gunaratne, David Critchley, Igor Barsukov, Edward Manser, Christoph Ballestrem

**Author notes:** Contributed equally (Paul Atherton, Rafaella Konstantinou).

## Abstract

The formation of healthy tissue involves continuous remodelling of the extracellular matrix (ECM). Whilst it is known that this requires integrin-associated cell-ECM adhesion sites (CMAs) and actomyosin-mediated forces, the underlying mechanisms remain unclear. Here we examine how tensin3 contributes to formation of fibrillar adhesions (FBs) and fibronectin fibrillogenesis. Using BioID mass spectrometry and a mitochondrial targeting assay, we establish that tensin3 associates with the mechanosensors talin and vinculin. We show that the talin R11 rod domain binds directly to a helical motif within the central intrinsically disordered region (IDR) of tensin3, whilst vinculin binds indirectly to tensin3 via talin. Using CRISPR knock-out cells in combination with defined tensin3 mutations, we show (i) that tensin3 is critical for formation of α5β1-integrin FBs and for fibronectin fibrillogenesis, and (ii) the talin/tensin3 interaction drives this process, with vinculin acting to potentiate it.

## Introduction

Cells interact with the extracellular matrix (ECM) through transmembrane integrin receptors that connect with the actin cytoskeleton through multi-protein complexes. Integrin-mediated cell-matrix adhesions (CMAs) can vary in location, shape and function. CMAs initially form at the cell periphery as dot-like focal complexes that develop into focal adhesions (FAs) through actomyosin-mediated tension. FAs can mature further into centrally located fibrillar adhesions (FBs). In contrast to the well-established roles of FAs in sensing both biophysical and biochemical ECM cues, rather little is known about the formation and function of FBs. FBs are intimately linked to fibronectin fibrils and other associated ECM proteins such as collagens, the organisation of which depends on integrins (Clark et al. 2005; Green et al. 2009; Pankov et al. 2000; Lu et al. 2020). The formation of fibronectin fibrils has been observed in both fibroblasts and epithelial cells (Lu et al. 2020), and involves the translocation of fibronectin-bound α5β1 integrins from peripheral FAs to central FBs (Clark et al. 2005; Pankov et al. 2000) in a tension-dependent manner (Zamir et al. 2000; Lu et al. 2020). The force-mediated maturation of FAs to FBs is accompanied by a switch in molecular composition: FAs are enriched in talin, vinculin, paxillin and FAK; FBs predominantly contain α5β1 integrins and tensin1 and 3 (Zamir et al. 2000; Clark et al. 2010). While overexpression of a dominant negative chicken tensin fragment and tensin depletion experiments show that tensins are important for fibronectin fibrillogenesis (Pankov et al. 2000; Georgiadou et al. 2017), mechanistic insights are lacking.

In contrast to the single tensin gene in chickens, mammals have four tensin genes (tensin1-4), three of which (tensin1-3) encode structurally similar proteins (Supp. Fig. 1a). Expression of fluorophore-tagged human tensin1-3 in fibroblasts revealed variations in localisation to different CMA types. Whilst tensin2 predominantly localised to FAs, and tensin3 to FBs, tensin1 was in both structures (Clark et al. 2010). What regulates this difference remains unclear given their structural similarities. Although several proteins have been found to bind to the highly conserved N- and C- termini of tensins (Liao and Lo 2021; Calderwood et al. 2003; Katz et al. 2007; McCleverty, Lin, and Liddington 2007; Zhao et al. 2016; Lo et al. 1994; Shih et al. 2015; Cao et al. 2012; Liao et al. 2007; Kawai et al. 2009; Qian et al. 2007; Cui, Liao, and Lo 2004; Muharram et al. 2014; Hall et al. 2010; Goreczny, Forsythe, and Turner 2018) (Supp. Fig. 1b), most of the interaction studies involved binding to isolated peptides, with little evidence that the interations occur in cells or are linked to specific biological functions.

In this study we aimed to gain mechanistic understanding of the role of tensins in FB formation and its contribution to fibronectin fibrillogenesis. We focussed on tensin3, the family member with strongest enrichment in FBs (Clark et al. 2010). Using a combination of proximity biotinylation mass spectrometry (BioID) and fluorescence microscopy, we identified several potential tensin interactors including the mechanosensors talin and vinculin. In vitro NMR and ITC experiments showed that the intrinsically disordered region (IDR) of tensin3 contains a talin binding motif similar to those previously identified in DLC1 and RIAM (Goult et al. 2013). Complementary cell biological assays showed that the tensin3-talin interaction is critical for force-mediated maturation of FBs and fibronectin fibrillogenesis, processes potentiated by vinculin binding to talin.

## Results

### Actomyosin-mediated forces are required for formation of FBs but not their maintenance

Immunostaining of U2OS osteosarcoma cells or Telomerase Immortalised Fibroblasts (TIF) with antibodies specific for the different tensin isoforms (for characterisation of antibody specificity see Supp. Fig. 1c), revealed that both cell types expressed tensin1 and tensin3, but little tensin2 (Supp. Fig. 1d). As reported for other cell types (Clark et al. 2010), tensin3 was enriched at centrally located CMAs, co-aligning with fibronectin fibrils, whereas vinculin was more prominent in peripheral adhesions (Fig. 1a, b, Supp. Fig. 1e). FBs were previously shown to develop from FAs via myosinII-dependent gliding of α5β1 integrins (Lu et al. 2020). We confirmed that a similar mechanism regulates tensin3 localisation by treating U2OS cells or TIFs with the actomyosin inhibitor blebbistatin (50 μM) prior to cell spreading on fibronectin. Under these conditions neither vinculin-positive FAs nor tensin3-enriched FBs formed. In contrast, when cells were treated with blebbistatin after spreading, tensin3-rich FBs remained but vinculin-positive FAs disappeared (Fig. 1c, Supp. Fig. 1f). These data not only confirm that FB formation involves myosinII contractility, but suggest that the formation of FAs and their associated molecular machinery are a prerequisite for FB development.

**Fig. 1.**
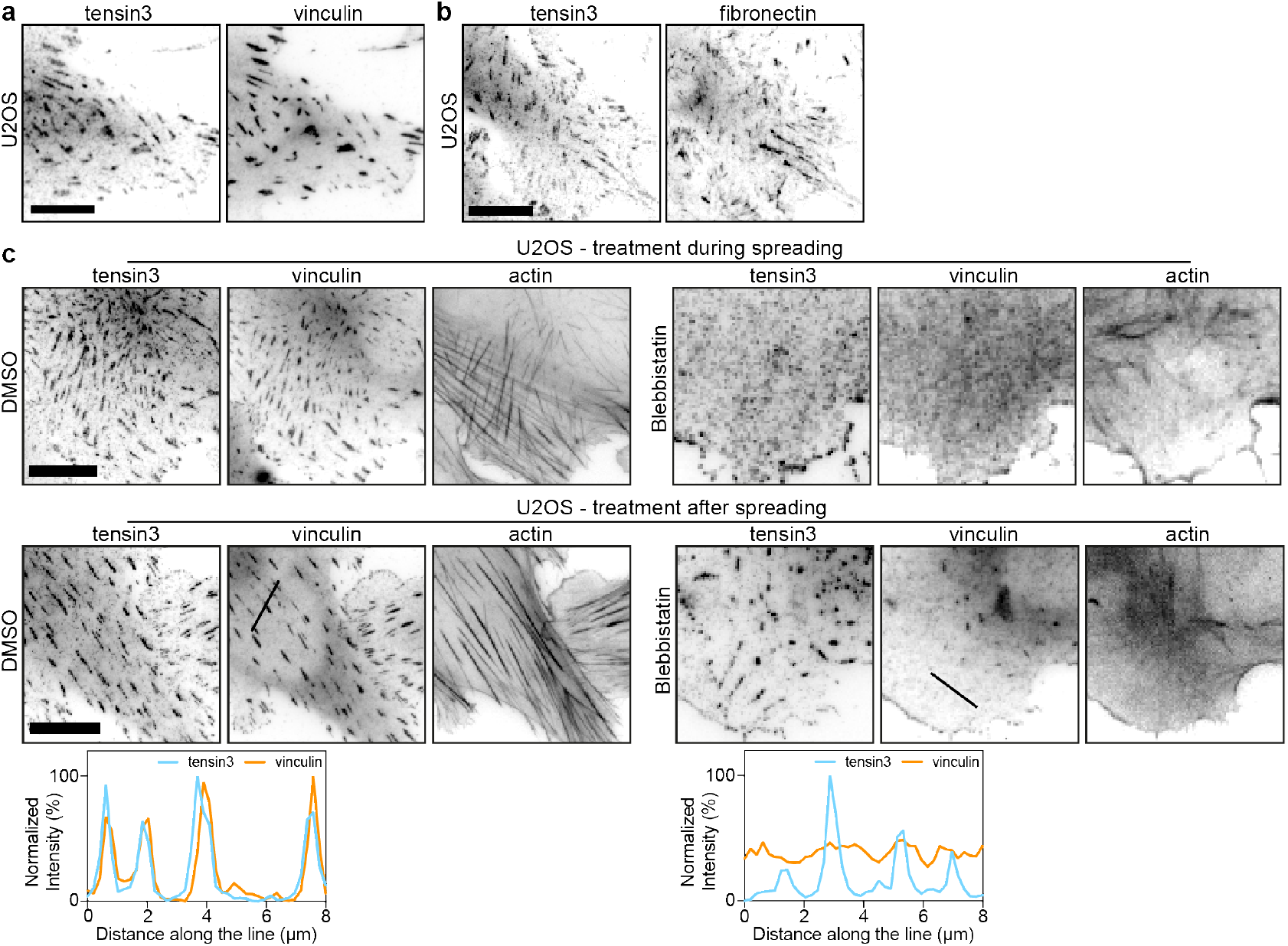
Actomyosin-mediated forces are required for the formation of tensin3-positive fibrillar adhesions. Representative images of immunofluorescence images of U2OS cells cultured overnight on fibronectin-coated glass stained for **(a)** tensin3 and vinculin, or **(b)** tensin3 and fibronectin. **(c)**, Upper panels: U2OS cells were treated in suspension with blebbistatin (50 μM) or an equivalent volume of DMSO, for 60 minutes. Cells were fixed after spreading on fibronectin-coated glass for 4 hours. Note the absence of tensin3- or vinculin-positive structures in blebbistatin-treated cells. Lower panels: U2OS cells cultured overnight on fibronectin-coated glass were treated with blebbistatin (50 μM) or an equivalent volume of DMSO, for 60 minutes prior to fixation. Line profiles indicate fluorescences intensity levels of proteins from the line profiles in the FA images from the lower panels in (c). Note that tensin3 remains at adhesions whilst vinculin disappears after blebbistatin treatment. Scale bars in a-c indicate 10 μm.

### Identification of tensin binding partners

To better understand the mechanisms underlying formation of tensin3-enriched central adhesions, we examined the molecular neighbourhood of tensin3 using SILAC (stable isotope labeling by amino acids) BioID which labels proteins within a 1—20 nm radius of the bait protein (Dong et al. 2016). We stably expressed tensin3 with an N-terminal BirA* plus myc tag in U2OS cells (BirA-tensin3); cells expressing myc-BirA* alone (BirA-control) were used as controls. Immunofluorescence showed that BirA-tensin3 localised prominently in FBs and a similar pattern was seen for biotinylation (Supp. Fig. 2a), indicating the construct is functional.

Mass spectrometry analysis using normalised heavy-to-light ratios as an index for molecular proximities with a threshold ≥2.5 identified 13 proteins in the 20 nm radius proximal to the N-terminal of tensin3 (Fig. 2a). Comparison with literature curated (Zaidel-Bar et al. 2007) and experimental datasets (Horton et al. 2015; Dong et al. 2016; Chastney et al. 2020) demonstrated that the majority of these tensin “neighbours” are known bona fide CMA proteins (Fig. 2b). Other FA proteins such as vinculin, LPP and kindlin-2, were found in a list of proteins with a heavy-to-light ratio threshold ≥1.0 (Supp. Table 1). Together, these data indicate that tensin3 is embedded in a dense network of CMA proteins. To validate a number of these potential interactions, and to extend our findings to tensin1 and 2, we performed experiments using a mitochondrial targeting and recruitment system (Atherton et al. 2020). For these assays tensin1-3 were tagged with mCherry at their N-termini, and with the cBAK mitochondrial targeting sequence (Atherton et al. 2020) at their C-termini. When expressed in NIH3T3 fibroblasts all of these tensin constructs localised to mitochondria (Supp. Fig. 2b). We then examined the ability of putative GFP-tagged tensin binding partners to co-localise with mCherry-tensin-cBAK at mitochondria (Fig. 2c, Supp. Fig. 2c). Interestingly, both tensin1 and tensin3 recruited talin (talin1 and 2), vinculin, paxillin, FAK, KANK2 and ILK to mitochondria, whereas tensin2 only recruited ILK and paxillin (Fig. 2d). Immunofluoresence confirmed the co-localization of endogenous talin1, vinculin and paxillin with tensin1 and 3 at mitochondria (Supp. Fig 2d). Integrins (active β1; 9EG7 staining) were absent from mitochondria (Supp. Fig. 2e), indicating that the associations occur independently of the reported tensin-integrin association (McCleverty, Lin, and Liddington 2007; Georgiadou et al. 2017).

**Fig. 2.**
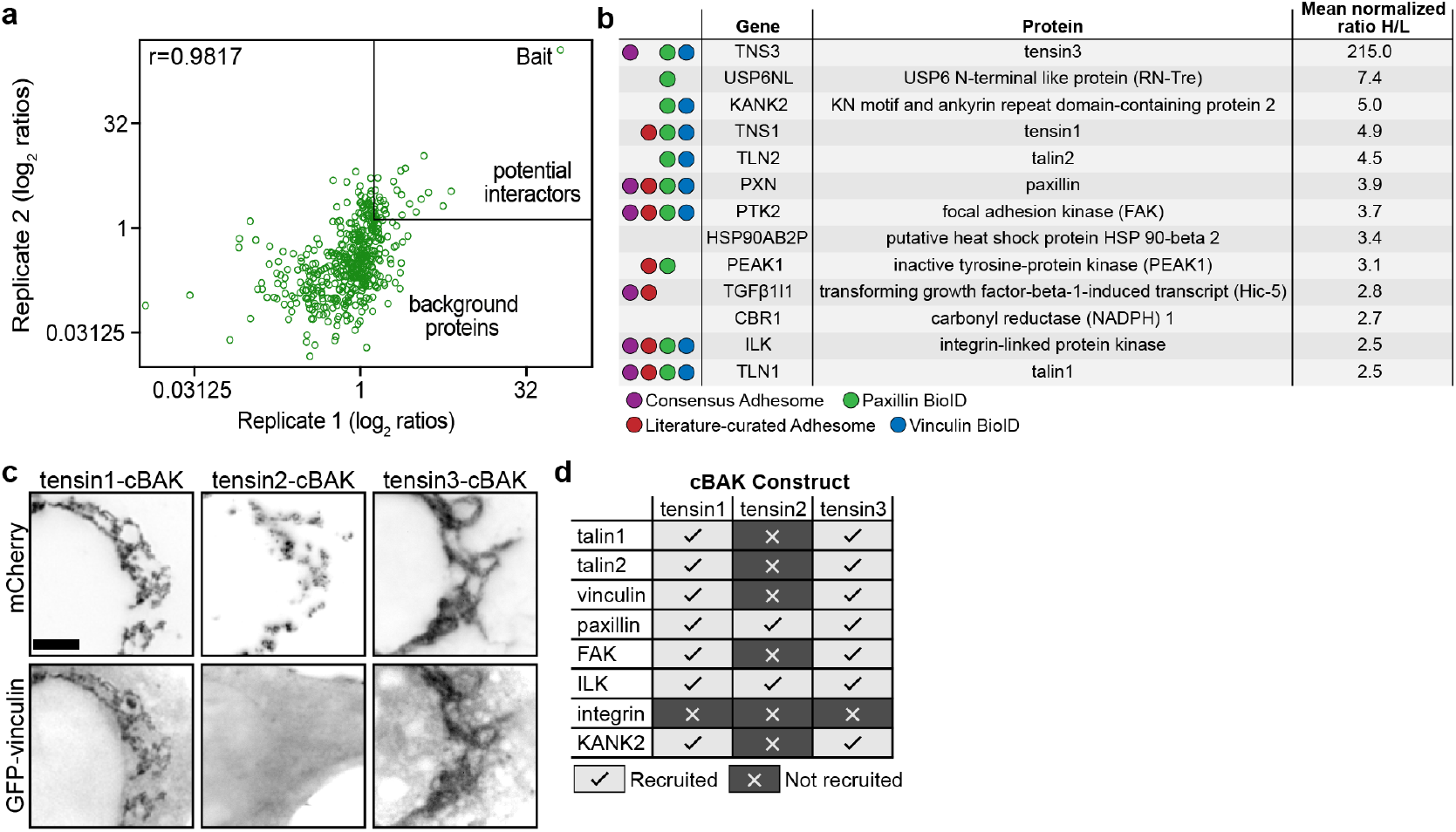
Tensin3 interaction partners revealed by proximity biotinylation (a, b) and the mitochondrial targeting system (c, d). **a**, Scatter plot showing the correlation of the SILAC normalised log2 ratio of the two tensin3 BioID repeats (data set in file S1). The majority of proteins are background proteins (bottom left of the graph); those on the top right of the graph (SILAC ratio *≥*2.5) were considered as enriched. The two repeats were highly reproducible with a correlation coefficient of r=0.9817. **b**, Table listing the proteins identified from the BioID experiments using a cut-off *≥*2.5 (Ratio of Heavy/Light). **c**, Co-expression of mCh-tensin1-, tensin2- or tensin3-cBAK with GFP-vinculin in NIH3T3 cells shows that GFP-vinculin co-localises at mitochondria with both tensin1- and tensin3-cBAK but not tensin2-cBAK. **d**, Summary table of which cell-matrix adhesion proteins co-localised with mCh-tensin1-, tensin2- or tensin3-cBAK in NIH3T3 cells; scale bar indicates 10 μm.

### Talin and vinculin associate with the tensin3-IDR

Following the BioID and mitochondrial targeting data plus our observations that FB formation is force-dependent (Fig. 1), we hypothesised a link between tensin and the mechanosensors talin and vinculin that connect integrins to the force exerting actin cytoskeleton (Atherton et al. 2015). To identify binding regions, we generated a series of tensin3 truncations (Fig. 3a, Supp. Fig. 3a) fused to GFP at the N-terminus. These included constructs lacking either the N- or C-terminal regions (‘ΔNterm’ and ‘ΔCterm’, respectively), and those consisting of the N- or C-terminal regions only (‘Nterm’ and ‘Cterm’). We first tested the recruitment of the GFP-tagged tensin truncations to mCh-vinFL-cBAK or mChtalin-cBAK expressed in NIH3T3 fibroblasts. Only tensin3 constructs containing the central IDR (GFP-tensin3ΔNterm and GFP-tensin3-ΔCterm) were recruited (Fig. 3b, c). This was surprising, since the N-terminal PTEN-like region and C-terminal SH2-PTB regions had previously been thought to target tensins to FA (Liao and Lo 2021). We therefore prepared constructs with the tensin3-IDR fused to cBAK or GFP and found that they associated with both talin and vinculin in mitochondrial targeting experiments (Fig. 3d, Supp. Fig. 3b). From these data we concluded that talin and vinculin associate directly or indirectly with the tensin3-IDR.

**Fig. 3.**
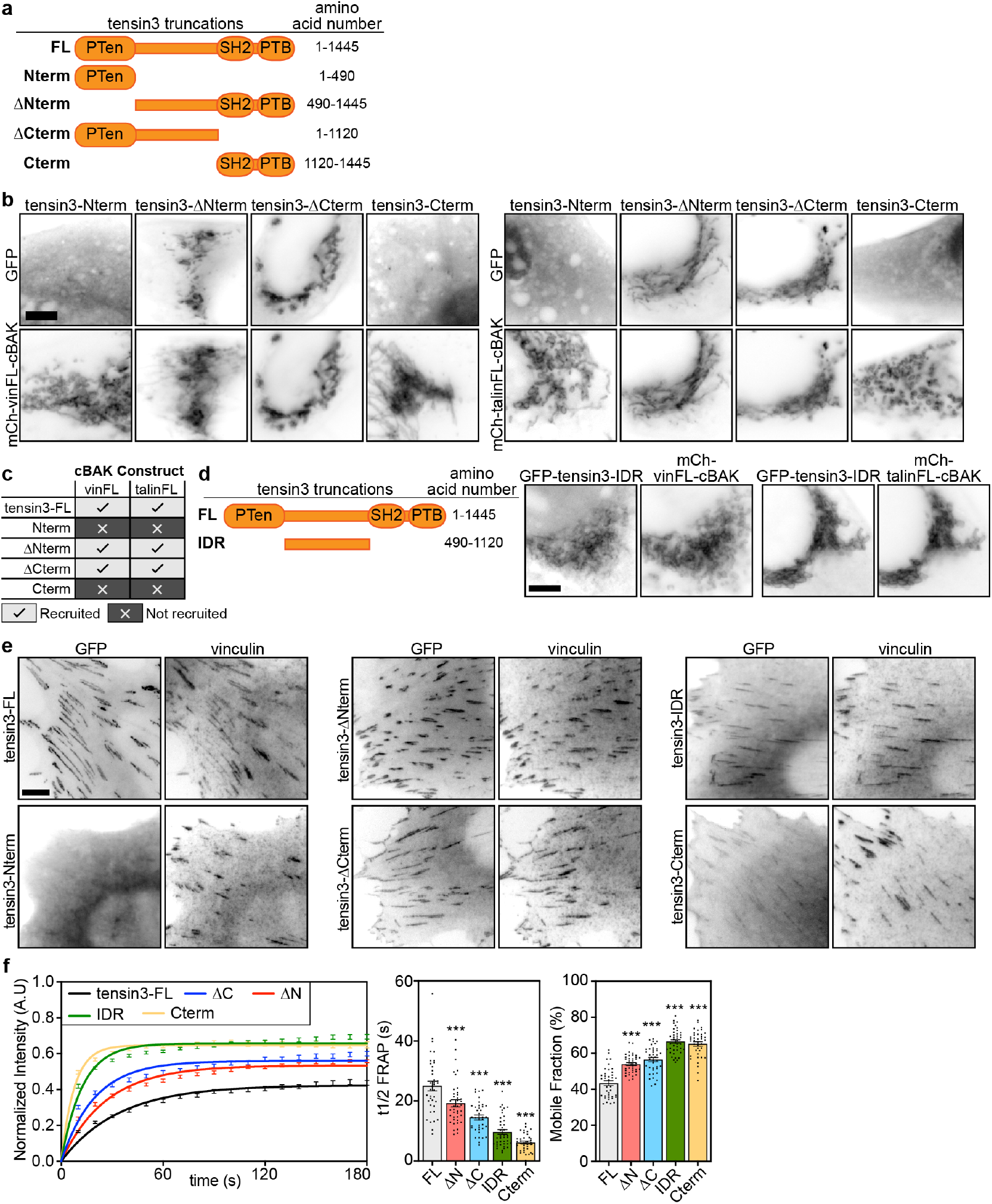
The tensin3 IDR is required for the interaction with talin and vinculin. **a**, Schematic of the tensin3 deletion mutants used, with indicated truncation sites. **b, c**, Tensin3 truncations were expressed as N-terminally tagged GFP fusion constructs together with mCh-vinFL-cBAK or mCh-talinFL-cBAK in NIH3T3 cells. Only those constructs containing the intrinsically disordered region (IDR) of tensin3 linking the PTen and SH2 domains co-localised with vinculin or talin at mitochondria. **d**, A tensin3 IDR-only construct (GFP-tensin3-IDR) co-localises at mitochondria when co-expressed in NIH3T3 cells with either mCh-vinFL-cBAK or mCh-talinFL-cBAK. **e**, GFP-tagged tensin3 truncations expressed in NIH3T3 cells; strongest localisation to cell-matrix adhesions is observed in those constructs containing the tensin3 IDR (GFP-tensin3-ΔNterm, GFP-tensin3-ΔCterm, GFP-tensin3-IDR) **f**, FRAP experiments in NIH3T3 cells expressing the indicated GFP tensin3 fusion constructs show that the tensin3-IDR contributes to tensin3 mobility within adhesion sites. Whereas GFP-tensin3-FL had a mean half-time of fluorescence recovery (t1/2 FRAP) of 25.00s ±1.56 and a mobile fraction of 43.34% ±1.36, both GFP-tensin3-ΔNterm and GFP-tensin3-ΔCterm showed significantly increased turnover (t1/2 FRAP 19.26s ±1.16 and 14.51s ±0.81 respectively; mobile fractions 53.99% ±0.80 and 56.46% ±1.15). Similarly, both the GFP-tensin3-Cterm and GFP-tensin3-IDR constructs had significantly increased turnover (t1/2 FRAP 6.16s ±0.48 and 9.66s ±0.81, respectively; mobile fractions 65.19% ±1.20 and 66.50% ±1.06). Error bars are S.E.M, n = 38 (GFP-tensin3-FL), 40 (GFP-tensin3-ΔNterm), 39 (GFP-tensin3-ΔCterm), 37 (GFP-tensin3-Cterm), 39 (GFP-tensin3-IDR) adhesions from 5 cells; *** indicates p<0.001 (One-way ANOVA) against tensin3-FL. Scale bars in b, d, e indicate 10 μm.

### The IDR contributes to tensin3 localisation to CMAs

The association of tensin3 with the FA proteins talin and vinculin through its IDR suggested that this region could contribute to the localisation of tensin3 to CMAs. To test this possibility, we expressed the GFP-tensin deletion mutants in NIH3T3 cells and assessed their co-localisation with endogenous vinculin. Both GFP-tensin3-ΔNterm and GFP-tensin3-ΔCterm strongly localised to CMAs (Fig. 3e). Albeit weaker, both the GFP-tensin3-Cterm and GFP-tensin3-IDR constructs localised to vinculin positive adhesions, whereas GFP-tensin3-Nterm localised to the cytoplasm (Fig. 3e). Comparing the dynamics of the deletion mutants with GFP-tensin3-FL in FRAP experiments revealed that the IDR contributes to tensin3 stability within CMAs (Fig. 3f; Supp. Fig. 3c). These data lead to the conclusion that both the central IDR and C-terminal SH2-PTB domains are critical for the recruitment of tensin3 to CMAs and for maximising its efficient binding to adhesion components.

### A helical motif in the tensin3-IDR binds talin

To date no structural domains or binding sites have been identified in the tensin-IDR that could account for binding to talin or any other FA protein. Examining the predicted secondary structure of the tensin3-IDR (Supp. Fig. 4a) identified two short amino acid sequences containing a combination of hydrophobic and charged residues predicted to form helices (helices H1, residues 690-710 and H2, residues 1030-1050 of tensin3; Fig. 4a). Mitochondrial targeting experiments using tensin3 constructs lacking H1 or H2 revealed H1 to be responsible for the interaction of tensin3 with talin and vinculin (Fig. 4b, c, Supp. Fig. 4b, c). We next tested the contribution of H1 to tensin3 CMA localisation. The noticeable increase of cytoplasmic fluorescence intensity in cells expressing a GFP-tensin3-FLΔH1 deletion mutant in comparison to GFP-tensin3-FL immediately suggested a reduced affinity for adhesion sites (Fig. 4d). This was confirmed by the significantly increased mobility of GFP-tensin3-FLΔH1 in comparison to GFP-tensin3-FL in FRAP experiments (Fig. 4e, Supp. Fig. 4d). However, the GFP-tensin3-FLΔH1 construct still localised to CMAs suggesting that additional regions of tensin3 also contribute to tensin localisation. Based on our previous results (Fig. 3), we hypothesised a role for the C-terminal SH2-PTB domains. We prepared constructs lacking H1 in either the tensin3-IDR (GFP-tensin3-IDRΔH1) or the tensin3-ΔCterm construct (GFP-tensin3-ΔH1ΔCterm). In contrast to GFP-tensin3-IDR or GFP-tensin3-ΔCterm, both of the ΔH1 constructs were absent from CMAs (compare Fig. 4f with Fig. 3e). From these results we conclude that both H1 and the C-terminus are the major sites that contribute to tensin recruitment to CMAs. Interestingly the H1 motif of tensin3 is conserved in the tensin1-IDR, but has very low homology with the tensin2-IDR (Fig. 4g), consistent with the similarity in tensin1 and tensin3 localisation (Supplementary Fig. 1). From these data we conclude that the H1 motif likely mediates the association of tensin3 with talin and vinculin and that this motif contributes to tensin3 localisation to CMAs.

**Fig. 4.**
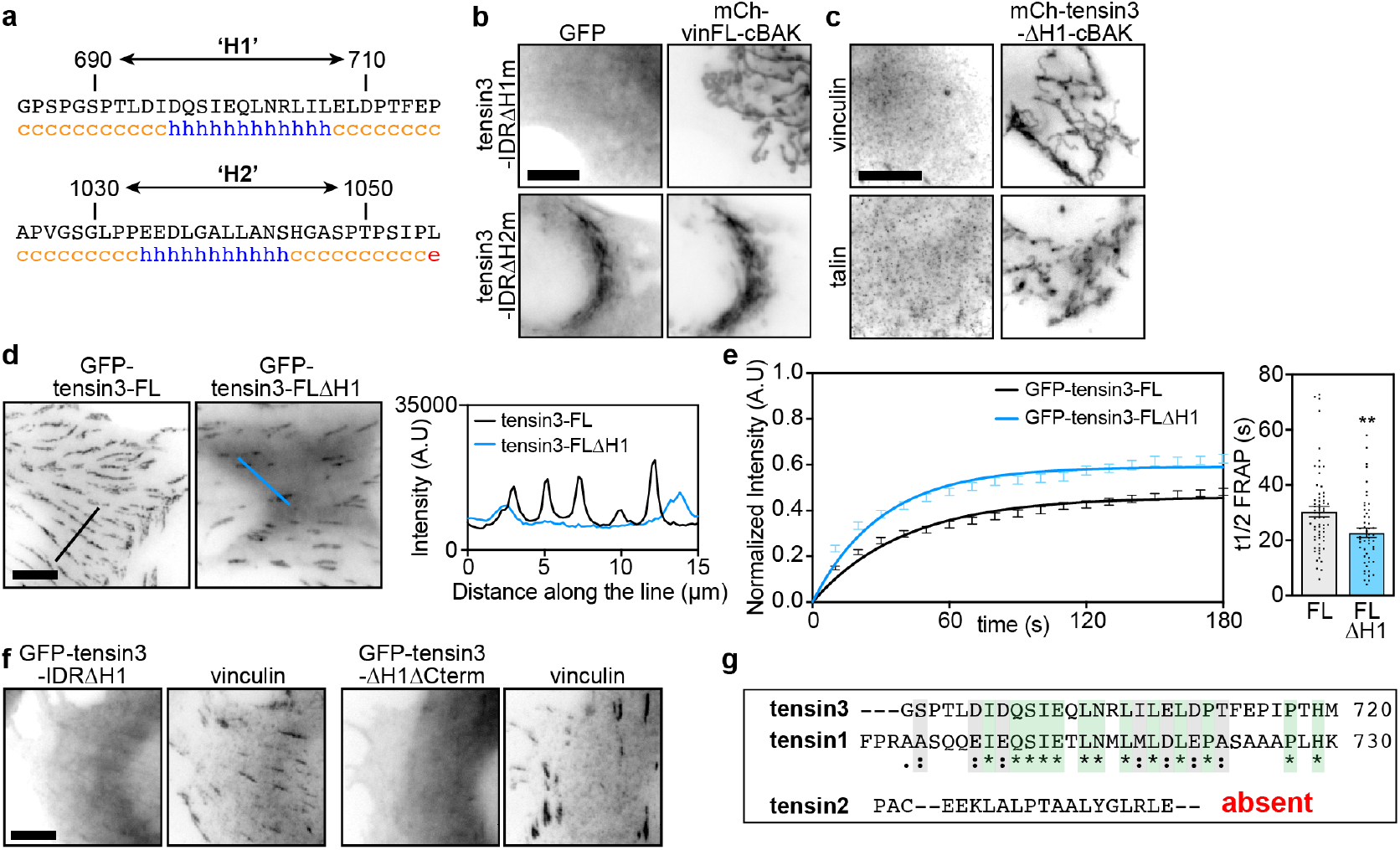
A helical motif in the tensin3 IDR mediates the association with talin and vinculin. **a**, Secondary structure prediction of the tensin3 IDR reveals two stretches of amino acids predicted to form a helix, termed ‘H1’ and ‘H2’. **b**, Co-expression of mCh-vinFL-cBAK together with a tensin3-IDR construct lacking either H1 (GFP-tensin3-IDRΔH1) or H2 (GFP-tensin3-IDRΔH2) in NIH3T3 cells shows that the H1 motif is responsible for the interaction between the tensin3 IDR and vinculin. **c**, Immunostaining of NIH3T3 cells expressing mCh-tensin3-ΔH1-cBAK shows that the H1 motif is responsible for the recruitment of endogenous vinculin and talin to mitochondria. **d**, A tensin3 construct lacking the H1 motif (GFP-tensin3-FLΔH1) expressed in NIH3T3 cells localises to cell-matrix adhesions. Line profiles show that this construct has a higher cytoplasmic fraction compared to GFP-tensin3-FL. **e**, FRAP experiments in NIH3T3 cells show the H1 motif regulates the turnover of tensin3 within cell-matrix adhesion sites; t1/2 FRAP (GFP-tensin3-FL and GFP-tensin3-FLΔH1) 30.40s ±1.85 and 22.76s ±1.73 respectively; mobile fraction 48.77% ±1.51 and 61.96% ±1.79. Error bars are S.E.M, n = 63 (GFP-tensin3-FL) or 51 (GFP-tensin3-FLΔH1) adhesions from 7 cells; *** indicates p<0.001 (t-test). **f**, Representative images of NIH3T3 cells expressing a tensin3 IDR construct lacking the H1 motif (GFP-tensin3-IDRΔH1) or a tensin3 construct lacking both the H1 motif and the C-terminal SH2-PTB domains (GFP-tensin3-ΔH1ΔCterm). Note that neither of these constructs shows localisation to vinculin-positive cell-matrix adhesion sites. **g**, Amino acid sequence alignment of tensin1, tensin2 and tensin3 shows that there is some homology between the tensin3 H1 region and a corresponding stretch of amino acids in tensin1. No homology is observed in tensin2. Scale bars in b, c, d, f indicate 10 μm.

### Tensin3 associates with vinculin indirectly through talin

Our finding that both talin and vinculin require the same H1 region in the IDR for association (Fig. 4c) raised the possibility that the association of tensin with one of them is direct, the other indirect;. To test this possibility, we performed mitochondrial targeting experiments in talin or vinculin knock out cells. Co-localisation of endogenous talin and mCh-tensin3-FL-cBAK expressed in vinculin knock-out MEFs clearly shows that the talin/tensin3 interaction is vinculin independent (Fig. 5a). In contrast, endogenous vinculin failed to associate with mCh-tensin3-FL-cBAK expressed in talin1 and 2 null cells (Fig. 5b). This clearly indicates that tensin association with vinculin requires at least one of the talin isoforms. From these results we hypothesised vinculin could indirectly regulate tensin3 function via talin. To explore this, we expressed GFP-tensin3-FL together with either mCherry-tagged wildtype (mCh-vinFL) or constitutively active (mCh-vinT12; (Cohen et al. 2005)) vinculin in vinKO MEFs, and measured adhesion formation after spreading on fibronectin in the presence of blebbistatin. Similar to our earlier observations in U2OS and TIF cells (Fig. 1), vinKO cells co-expressing GFP-tensin3-FL and mCh-vinFL formed few GFP-tensin3-FL-positive adhesions in the absence of actomyosin-mediated tension. In contrast, vinKO cells co-expressing mCh-vinT12 and GFP-tensin3-FL displayed large GFP-tensin3-FL positive adhesions. Importantly, this effect was not seen in cells co-expressing mCh-vinT12 and the GFP-tensin3-FLΔH1 mutant unable to bind talin (Supplemental Fig. 5a). Neither was this effect seen in cells expressing an active vinculin construct unable to bind talin (vinT12-A50I) (Bakolitsa et al. 2004) (Supp. Fig. 5b), supporting the notion that vinculin regulates tensin3 indirectly via talin.

**Fig. 5.**
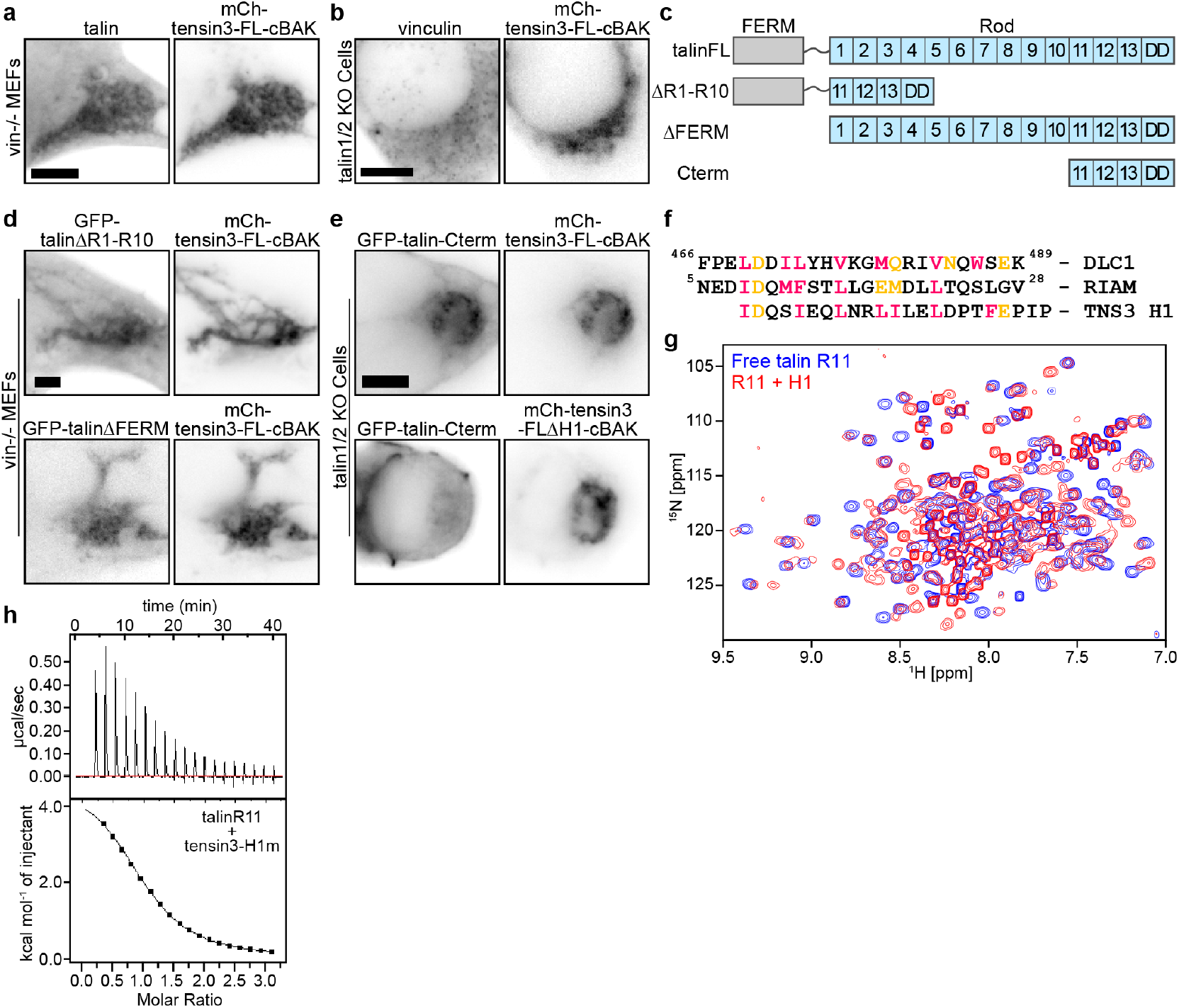
Tensin3 H1 binds directly to the R11 domain of talin. Immunostaining of **(a)** endogenous talin in vinculinKO mouse embryonic fibroblasts (vin-/-MEFs) or **(b)** endogenous vinculin in talin1/2 KO cells expressing mCh-tensin3-FL-cBAK reveals that vinculin interacts with tensin3 indirectly through talin. **c**, Schematic of talin constructs used. **d**, Representative images of vin-/- MEFs cells expressing GFP-talinΔR1-R10 or GFP-talinΔFERM together with mCh-tensin3-FL-cBAK. **e**, Representative images of talin1/2 KO cells expressing GFP-talin-Cterm with either mCh-tensin3-FL-cBAK or mCh-tensin3-FLΔH1-cBAK. **f**, Sequence alignment of the talin binding sites in DLC1 and RIAM with the H1 sequence of TNS3. **g**, 1H,15N-HSQC NMR spectra of 400 μM 15N-labelled isolated talin R11 in the absence (blue) and presence (red) of 1,600 μM of tensin H1 peptide. Note the decrease of signals in the complex due to the exchange broadening. **h**, ITC measurements of tensin3 H1 interactions with talin R11. The experiments were conducted with 80 μM of R11 in cell and 1.2 mM of the peptide in the syringe. Scale bars in a, b, d, e indicate 5 μm.

### Tensin H1 binds to the talin R11 rod domain

To identify the region in talin that binds the tensin H1 motif, we co-expressed a variety of talin deletion constructs (Fig. 5c) in vinculin knock-out MEFs together with mCh-tensin3-FL-cBAK. Initially, we analysed the recruitment of either a ΔFERM domain talin construct (GFP-talinΔFERM), or a talin construct lacking the R1-R10 rod domains (GFP-talinΔR1-R10) to mitochondria. Both talin constructs localised with mCh-tensin3-FL-cBAK at mitochondria (Fig. 5d) implicating the C-terminal talin R11-R13 rod domains in the interaction. Indeed, experiments in talin1/2 KO cells confirmed that a talin R11-R13 construct (GFP-talin-Cterm) colocalised with mCh-tensin3-FL-cBAK at mitochondria (Fig. 5e). Importantly, this GFP-talin-Cterm construct did not co-localise with mCh-tensin3-FLΔH1-cBAK (Fig. 5e), confirming that it binds to the tensin IDR H1 motif.

To test this interaction at CMAs, we co-expressed the GFP-talin-Cterm construct together with either mCh-tensin3-FL or mCh-tensin3-FLΔH1 in talin1/2 KO cells. These cells only spread on fibronectin in the presence of ^Mn2+^; in line with our mitochondrial targeting results, cells expressing GFP-talin-Cterm and mCh-tensin3-FL formed adhesions, whereas cells expressing GFP-talin-Cterm and mCh-tensin3-FLΔH1 formed only small dot-like complexes (Supp. Fig. 5c).

Interestingly, the H1 motif shares similarities to the talin binding sites in RIAM and DLC1 (Fig. 5f) (Zacharchenko et al. 2016; Goult et al. 2013), suggesting that tensin may interact with the R11 domain of talin that incorporates a RIAM binding site (Goult et al. 2013). To test this hypothesis, we conducted NMR and ITC experiments using a synthetic peptide corresponding to tensin3 H1 (residues 692-718) and a recombinant talin R11 domain. Addition of the tensin3 H1 peptide to 15N-labelled talin R11 resulted in large chemical shift changes and broadening (Fig. 5g), clearly indicating an interaction between talin R11 and the tensin3 H1 peptide with a dissociation constant (Kd) in the μM range. The ITC data supported the conclusions from the NMR experiment, showing an endothermic interaction with a Kd of 17 μM and enthalpy of 4.8 kcal/mol (Fig. 5h). Together these data reveal that H1 motif in the tensin3-IDR is a bona fide talin binding site (TBS) that interacts with the talin R11 domain with an affinity similar to other reported T BSs. From hereon in we refer to the H1 motif as “TBS”.

### Tensin controls integrin activity in centrally located CMAs

Both talin and tensin bind and activate integrins (Calderwood, Campbell, and Critchley 2013; McCleverty, Lin, and Liddington 2007; Georgiadou et al. 2017). However, whilst talin is absolutely critical for the formation of peripheral FAs and cell spreading through integrin activation (Zhang et al. 2008; Atherton et al. 2015), tensin seems less important for cell spreading, but is important for fibronectin fibrillogenesis (Pankov et al. 2000). In our experiments, CRISPR mediated tensin3 knock-out in U2OS cells (Supp. Fig. 6a) confirmed this notion, as cells without tensin3 were still able to spread, but had dramatically reduced α5β1 integrin positive CMAs, which were located predominantly at the cell periphery (Fig. 6a, Supp. Fig. 6b). Similar results were seen by depleting tensin3 using siRNA in TIFs (Supp. Fig 6c, d). Interestingly, overexpression of GFP-tensin3-FL, but not GFP-tensin3-FLΔTBS, markedly increased the number of α5β1 integrin positive CMAs when expressed in U2OS TNS3 KO cells (Fig. 6b). Similar results were seen in talinKO cells co-expressing GFP-talin with either mCh-tensin3-FL or mCh-tensin3-FLΔTBS, with a striking absence of centrally-located CMAs in cells expressing mCh-tensin3-FLΔTBS (Fig. 6c). It is thus the expression of tensin3 and the co-operation with talin that seem to regulate the abundance of FBs.

**Fig. 6.**
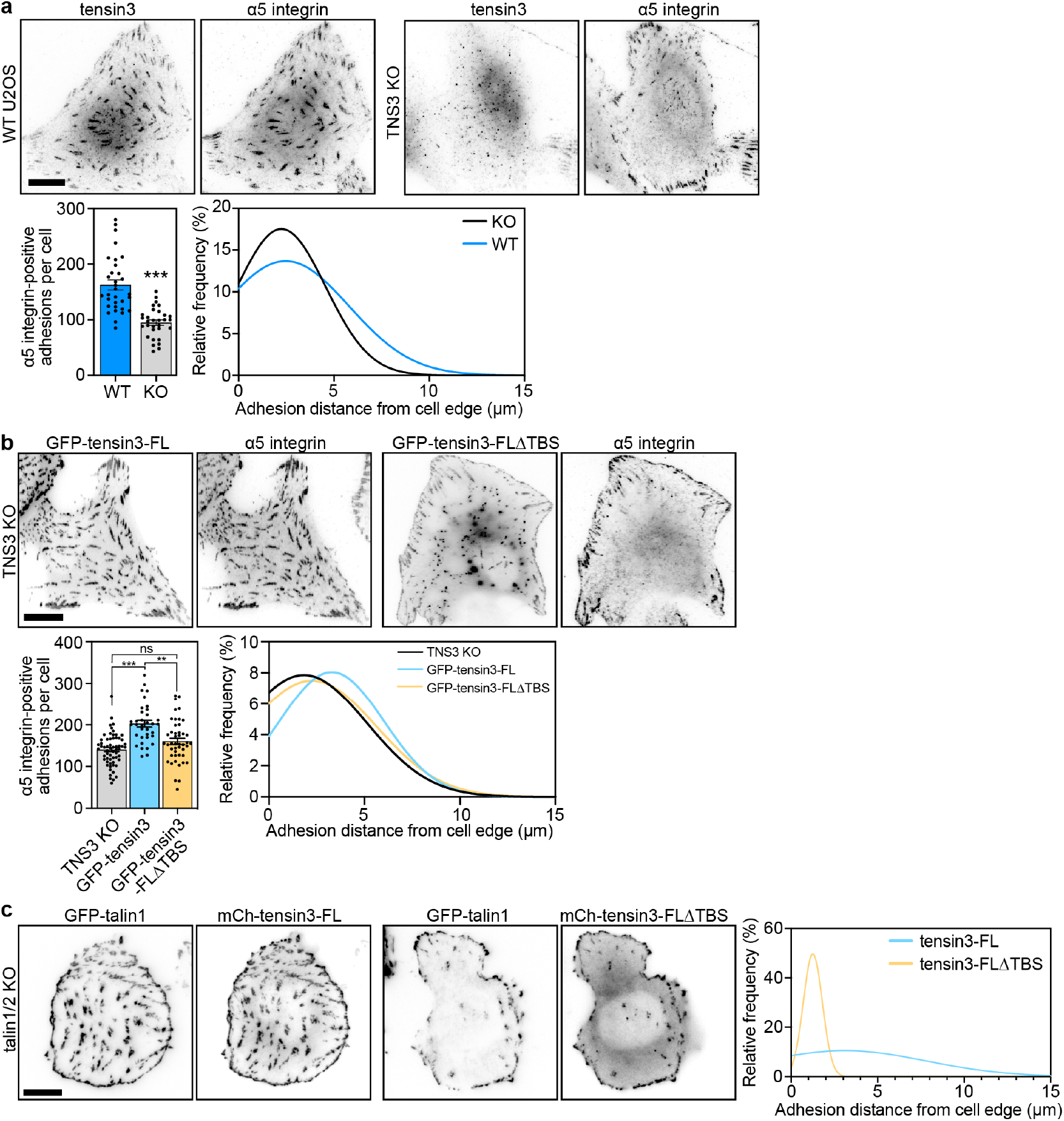
The tensin3 talin binding site regulates α5 integrin localisation. **a**, Representative images of wild-type (WT) or CRISPR-mediated TNS3 KO (TNS3 KO) U2OS cells spread overnight on fibronectin-coated glass prior to fixation and immunostaining for α5 integrin. Quantification shows that tensin3 KO cells (clone #2) have reduced α5 integrin-positive adhesions; error bars are S.E.M, n = 32 (WT) and 31 (KO) cells; *** indicates p<0.001 (t-test). Gaussian distribution of the distance of all detected α5 integrin-positive structures from the cell periphery shows that TNS3 KO cells have a reduced number of centrally-located adhesions. **b**, Representative images of TNS3 KO cells expressing the indicated GFP-tagged tensin3 construct fixed and stained for α5 integrin. Number of adhesions and the distance from the cell periphery was quantified as above, revealing that expression of the GFP-tensin3-FLΔTBS construct is unable to rescue the formation of centrally-located α5 integrin-positive adhesion structures. Error bars are S.E.M, n = 61 (TNS3 KO), 35 (GFP-tensin3), 46 (GFP-tensin3-FLΔTBS) cells; ** indicates p<0.01, *** indicates p<0.001 (ANOVA). Results in (a) and (b) are pooled from three independent repeats. **c**, Representative images of talin1/2 KO cells co-expressing GFP-talin1 and either mCh-tensin3-FL or mCh-tensin3-FLΔTBS. Co-expression of GFP-talin and mCh-tensin3 dramatically increased CMA formation and in particular the fraction of central adhesions. Graph shows the Gaussian distribution of the adhesion distance from the cell periphery; n = 19 (mCh-tensin3-FL) or 22 (mCh-tensin3-FLΔTBS) cells. Scale bars in a-c indicate 10 μm.

### The talin-tensin3 interaction regulates fibronectin fibril-logenesis

Given that tensin3 is required for the maturation of centrally-located α5-integrin positive adhesions, we next tested the ability of cells lacking tensin3 to produce fibronectin fibrils. Strikingly, fibronectin fibrillogenesis (assessed after culture overnight using an antibody against cell-derived fibronectin (Serini et al. 1998)) was reduced by approximately 50% in U2OS TNS3 KO cells compared to WT (Fig. 7a), with similar results observed in TIFs after siRNA-mediated knockdown of tensin3 (Supp. Fig. 7a). From these results, we hypothesised that cells expressing tensin3 lacking the IDR TBS would also show impaired fibronectin fibrillogenesis. Whereas expression of GFP-tensin3-FL in U2OS TNS3 KO cells rescued fibronectin fibrillogenesis to similar levels seen in WT U2OS cells, expression of GFP-tensin3-FLΔTBS had little/no effect (Fig. 7b). These data clearly demonstrate that the TBS in tensin3 is critically involved in fibronectin fibrillogenesis.

**Fig. 7.**
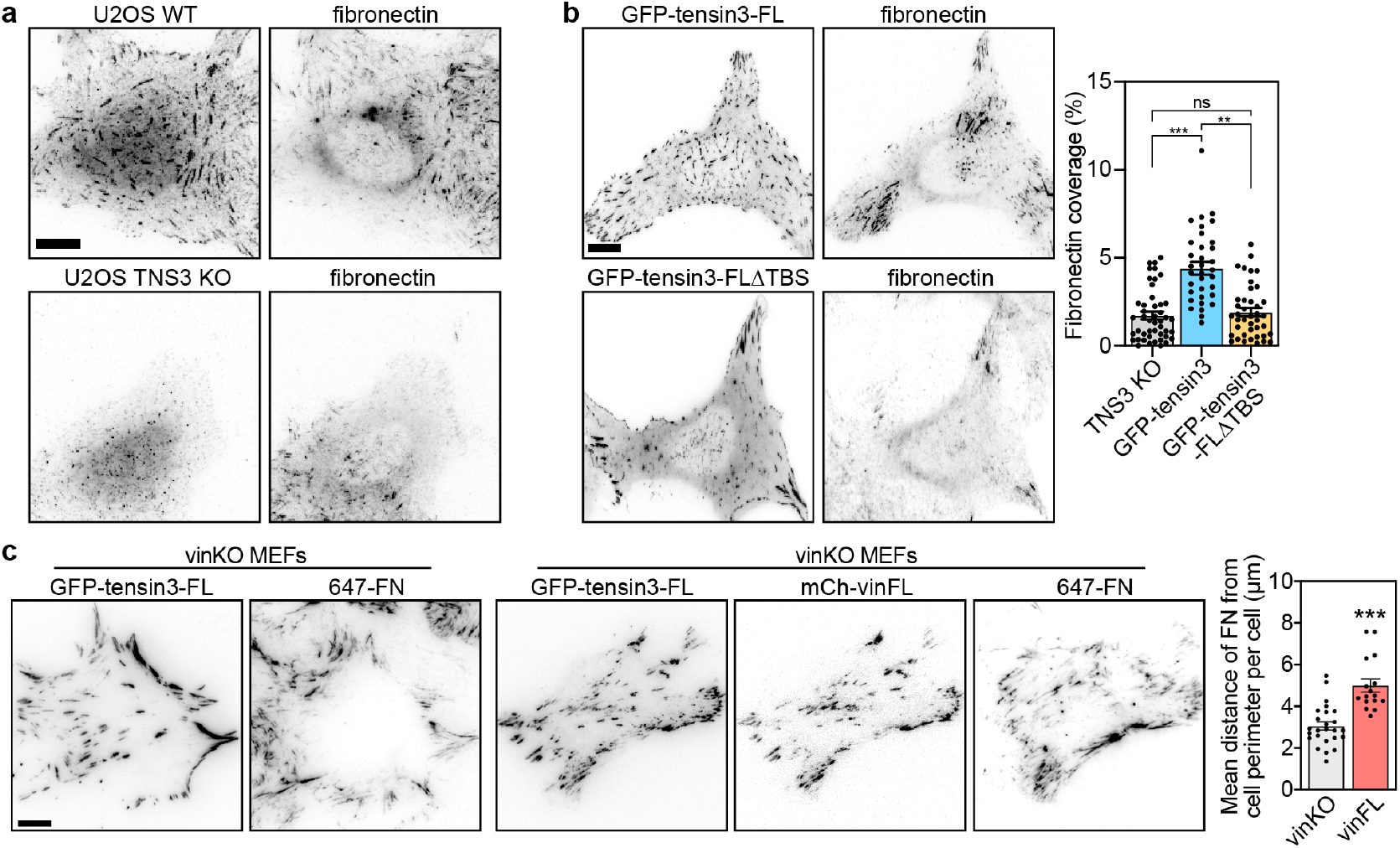
Fibronectin fibrillogenesis is dependent on the tensin3-talin interaction. Representative images of **(a)** WT or TNS3 KO U2OS cells or **(b)**TNS3 KO cells expressing GFP-tensin3-FL or GFP-tensin3-FLΔTBS spread overnight on fibronectin-coated glass immunostained for fibronectin. Quantification of the cell area containing fibronectin fibrils shows that TNS3 KO cells produce fewer fibronectin fibrils compared to cells rescued with GFP-tensin3-FL. Note that expression of GFP-tensin3-FLΔTBS is unable to rescue fibronectin fibril formation. Error bars are S.E.M, n = 46 (TNS3 KO), 35 (GFP-tensin3), 41 (GFP-tensin3-FLΔTBS) cells; ** indicates p<0.01, *** indicates p<0.001 (ANOVA). Data are pooled from three independent repeats. **c**, Representative images of vinKO MEFs expressing GFP-tensin3-FL with or without mCh-vinFL. Alexa Fluor 647-labelled fibronectin (647-FN) was added to the culture medium for 2 hours prior to fixation. Quantification of the mean distance of fibronectin (FN) fibres from the cell periphery shows that cells without vinculin have significantly fewer centrally-located FN fibrils compared to cells expressing mCh-vinFL. Error bars are S.E.M; n = 25 (vinKO) or 16 (+vinFL) cells; *** indicates p<0.001 (t-test). Scale bars in a-d indicate 10 μm.

### Vinculin regulates tensin dynamics to control fibronectin fibrillogenesis

The above experiments demonstrate that the talin/tensin interaction is key for efficient FB formation and fibronectin fibrillogenesis. Vinculin is a critical regulator of talin function and stability at CMAs (Atherton et al., 2015); therefore we explored the contribution of vinculin to the regulation of tensin3 at CMAs by analysing adhesion dynamics using live-cell imaging of vinKO MEFs expressing GFP-tensin3-FL with or without mCh-vinFL. The absence of vinculin increased the mean speed of GFP-tensin3-FL adhesions by approximately 40% and reduced adhesion lifetime by approximately 20% (Supp. Fig. 7B, Supp. Movie 1). Since tensin and α5-integrin dynamics are thought to be involved in fibronectin fibrillogenesis (Pankov et al., 2000) we hypothesised that the reduced stability of tensin3-positive adhesions in vinculinKO MEFs could translate to impaired fibronectin fibrillogenesis. To assess this we added fluorescently-labelled fibronectin to the culture medium of vinKO MEFs expressing GFP-tensin3 with or without mCh-vinFL and quantified fibril formation after 2 hours. Whereas both cells had fibronectin fibrils at the cell periphery, centrally-located fibrils were largely absent from cells without vinculin (Fig. 7C, Supp. Fig. 7C). Taken together, our results lead to the conclusion that tensin3 associates with talin through the TBS in the tensin3 IDR, with vinculin functioning as an “enhancer” of the talin-tensin mediated adhesion maturation that is required for efficient fibronectin fibrillogenesis.

## Discussion

The cells’ ability to remodel their extracellular matrix is critical for tissue homeostasis. CMAs are known to play a key role in matrix organisation and FBs are thought to have a major role in this process (Clark et al. 2005). Actomyosin-mediated forces drive the maturation of FBs from FAs, a transformation associated with the enrichment of tensin3 alongside α5β1 integrin, both of which co-localise with fibronectin fibrils. Here we shed light on the mechanisms regulating this phenomenon. We demonstrate that FB maturation and fibronectin fibrillogenesis is controlled by the interaction of the talin R11 rod domain with a helical TBS motif in the tensin3 IDR (Fig. 5). Abolishing this interaction by depleting cells of tensin3, or through deletion of the tensin3 TBS, inhibits both α5β1 enrichment at FBs and fibronectin fibrillogenesis (Figs. 6, 7, Supp. Fig. 6, Supp. Fig. 7). Furthermore, we show that vinculin associates with tensin3 indirectly through talin, and acts to potentiate the tensin3-dependent FB maturation (Fig. 7) that drives fibronectin fibrillogenesis.

Our BioID data identified talin as a close tensin3 neighbour, providing initial clues about a possible interaction between tensin3 and the mechanosensory talin/vinculin complex. Whilst the proximity of tensin3 to both talin1 and 2 and KANK2 were novel findings, p revious B ioID experiments with FAK, vinculin, LPP, ILK, parvin (Chastney et al. 2020), paxillin and kindlin2 (Dong et al. 2016) as bait identified tensin3 as prey. How these multiple potential interactions with tensin3 are organised in time and space, and to what extent they collaborate to regulate tensin-associated functions remains to be determined. However, it is interesting to note that a KANK2/talin1 complex was found to have a role in the gliding of KANK2-positive CMAs into α5β1 enriched adhesions that may be similar to FBs (Sun et al. 2016).

The mitochondrial targeting system confirmed the BioID outcomes, and demonstrated that many of the identified neighbours form a complex with tensin3 independent of integrins which are absent from mitochondria (Fig. 2, Supp. Fig. 2). The overall similarity in (i) tensin1 and 3 binding partners and (ii) their localisation suggests that they have a similar function. However, the observation that tensin3 depletion leads to an almost complete loss of central adhesions demonstrates that tensin1 cannot compensate for loss of tensin3. In contrast, our results for tensin2 were quite different: whilst all tested binding partners for tensin3 also associated with tensin1, only a few including paxillin and ILK bound to tensin2 suggesting that tensin2 has a quite distinct function. Force-dependent maturation of peripheral FAs to central FBs was observed in early studies characterising FBs as structures enriched in tensin, fibronectin and α5β1 integrins (Pankov et al. 2000; Zamir et al. 2000). Our finding that tensin3 constructs lacking either the reported N-terminal actin binding site or C-terminal integrin binding site localised to central adhesions similar to tensin3 wild-type (Fig. 3e) suggest neither interaction drives force-dependent enrichment of tensin3 at FBs. Rather, our data reveal that it is the newly-identified tensin3-talin interaction (Fig. 5) that drives α5β1 positioning (Fig. 6) and fibronectin fibillogenesis (Fig. 7). FB formation was previously shown to be increased on stiffer substrates (Barber-Perez et al. 2020), where forces acting on talin are higher (Kumar et al. 2016) and talin turnover is reduced (Stutchbury et al. 2017). The finding that vinculin-null fibroblasts had fewer centrally-located fibronectin fibrils (Fig. 7d) and more dynamic tensin3-positive CMAs (Supp. Fig. 7c) suggests vinculin mediates the forces acting on the tensin3-talin complex which govern α5β1 positioning and fibronectin fibillogenesis as FAs transition to FBs. This is consistent with a previous study showing vinculin is required for force transmission and efficient ECM remodelling in 3D culture (Thievessen et al. 2015). Taken together, our findings suggest a new model (Fig. 8) which explains how tensin3-enriched FBs form and how they are involved in ECM re-modelling.

**Fig. 8.**
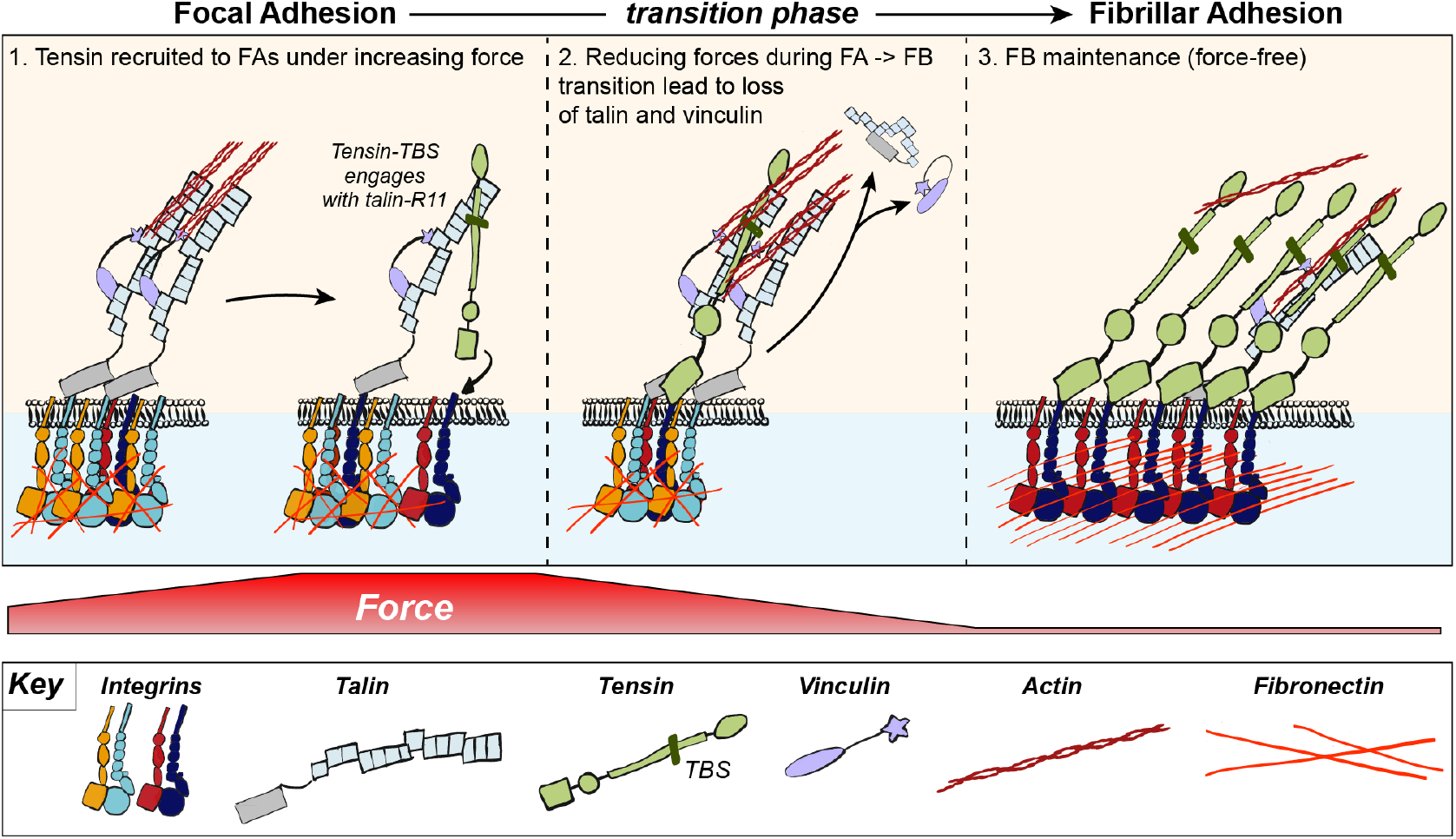
Model of tensin3 recruitment to CMAs. At the cell periphery actomyosin links integrins to adhesion complexes through talin and vinculin to form FAs. Further maturation of FA to FBs requires tensin binding to talin via the talin binding site located in the tensin IDR which engages the R11 domain of the talin rod. Binding of tensin3 to talin facilitates association of the C-terminal tensin3 PTB domain with the cytoplasmic tails of α5β1 integrins. FAs under forces and associated with fibronectin fibrils transition to form FBs that slide towards more central areas of cells. In these areas forces acting on the adhesion proteins decrease, leading to the release of the mechanosensors talin and vinculin. However, tensin3 remains present at FBs maintaining α5β1 integrins in an active, ligand-engaged state (Georgiadou et al. 2017) independently of tension. These stable integrin-tensin enriched structures remain attached to fibronectin fibrils and represent a signalling hub that remains active to uphold tension-independent integrin-mediated cell-matrix communication processes required to maintain tissue function.

## ACKNOWLEDGEMENTS

We thank the staff of the Bioimaging facility at the University of Manchester (in particular Dr. Peter March) for their assistance with microscopy. C.B. acknowledges the Biotehnology and Biological Sciences Research Council (BBSRC) and the Wellcome Trust for funding of this project. The Ballestrem laboratory is part of the Wellcome Trust Centre for Cell-Matrix Research, University of Manchester, which is supported by core funding from the Wellcome Trust (grant number 203128/Z/16/Z). P. Atherton was funded by BBSRC (BB/P000681/1 and BB/V016326/1); R. Konstantinou is supported by the Faculty of Biology, Medicine and Health and the University of Manchester and by the Agency of Science Technology and Research (A*STAR); E. Wang is funded by the BBSRC doctoral training programme in Liverpool. The Bioimaging Facility microscopes were purchased with grants from the BBSRC, the Wellcome Trust, and the University of Manchester Strategic Fund.

## AUTHOR CONTRIBUTIONS

**P.A.** Conceptualization, supervision, data curation, formal analysis, investigation, visualization, project administration, writing (original draft; review & editing); **R.K.** Investigation, data curation, formal analysis, visualization, writing (original draft); S.P.N. data curation, formal analysis; **E.W.** Investigation; **E.B.** Investigation; **M.P.**, **H.B.**, **K.C.** Resources; **J.G.** Data curation, writing (review); **D.C.** Writing (review & editing); **I.B.** Supervision, writing (review & editing); **E.M.** Supervision, project administration, writing (review editing); **C.B.** Conceptualization, supervision, funding acquisition, project administration, writing (original draft; review & editing).

## COMPETING FINANCIAL INTERESTS

The authors declare no competing financial interests.

## Materials and Methods

### Cell Culture

NIH3T3s, TIFs, U2OS and vinculin-/- MEFs were cultured in Dulbecco’s modified Eagles medium (DMEM) supplemented with 10% FCS (Lonza), 1% L-glutamine (Sigma), and 1% Non-essential amino acids (Sigma). Talin1&2 double null cells (Atherton et al. 2015) were cultured in DMEM:F12 (Lonza) supplemented with 10% FCS (Lonza), 1% L-glutamine (Sigma), 15 μM HEPES (Sigma) and 1% Non-essential amino acids (Sigma). Transient transfections were performed using Lipofectamine 2000 and Lipofectamine Plus reagents (Invitrogen), as per the manufacturer’s instructions. siRNA sequences used for transient knockdown of human tensin3 are given in Supp. Table 2. For live-cell imaging and fixed cell imaging, cells were cultured on glass-bottom dishes (IBL, Germany) coated with bovine fibronectin (Sigma) at a final concentration of 10 μg ml^−1^.

### Antibodies and Reagents

Samples were fixed in 4% paraformaldehyde (PFA), warmed to 37°C, for 15 minutes before being washed thrice with PBS. For immunofluorescence, samples were permeabilized at room temperature with Triton X-100 (0.5%) for 5 minutes, before being washed thrice. Primary and secondary antibodies used are given in Supp. Table 2. Blebbistatin (Tocris Bioscience) was diluted in DMSO (Sigma) and used at a final concentration of 50 μM. Mitotracker Deep Red FM (Thermo Fisher) was dissolved in DMSO to a concentration of 1 mM. Prior to use, the stock was diluted in pre-warmed medium at a final concentration of 200 nM, before being added directly to cells 30 minutes prior to fixation. Site-directed mutagenesis was performed using the QuikChange Lightning site-directed mutagenesis kit (Agilent) according to the manufacturer’s instructions.

### Protein extraction and western blot

Cells from a 6-well plate were lysed in 150 μl lysis buffer (25 mM HEPES pH 7.3, 150mM NaCl, 5 mM MgCl2, 1 mM EDTA, 20mM β-glycerophosphate, 5% glycerol, 0.5% Triton-X100, protease inhibitor). 20 μg of protein were loaded on 7.5% SDS-PAGE gel. The gel was transferred to a nitrocellulose membrane which was blocked in 5% milk. The membrane was probed for anti-tensin1, anti-tensin2, anti-tensin3, anti-GAPDH or anti-GFP. Primary antibody signal was detected using HRP-conjugated secondary antibodies imaged with an Azure c400 imaging system (Azure Biosystems), or using IRDye–conjugated secondary antibodies (LI-COR Biosciences) imaged with an Odyssey imaging system (LI-COR Biosciences).

### Plasmid preparation

The BioID vector pCDH-TagBFP-T2A-myc-BirA* (generous donation from Andrew Gilmore’s lab) was linearized (2 μg) by cutting with Xhol and BamH1. Gibson Assembly was used to design the primers to replace Xhol to BamHI region of the vector backbone with tensin3 (fragment 1: 2228 bp, forward primer: tggatgggcggagaaatctccctgagaagctcgGGTGGGTCCGGCGGTGGCTCTG-GCatggaggagggccatgg, reverse primer: ccgagggcgttcatgtgggtagggatgggctc; fragment 2: 2246 bp, forward primer: caccttcgagcccatccctacccacatgaacg, reverse primer: TCGACTCAGCGGTTTAAACTTA AGCTTGGTACCGAGCTCGtca-gaccttctttggtgaaccaatcatgacc). The two tensin fragments were amplified individually by PCR using KOD Xtreme™ Hot Start DNA Polymerase (Sigma) and were run on 1% agarose gel along with the linearized vector. Bands of correct size were cut out and purified using the Isolate II PCR and gel kit (Bioline). 50 ng of vector and 150 ng of each fragment were mixed with 3 μl of HiFi DNA Master Mix (NEB) and incubated at 50°C for 1 hour to form the new pCDH-TagBFP-T2A-myc-BirA*-Tensin3 construct. Bacterial transformation was then carried out using C2987 cells with 250 μl SOC medium. The DNA was extracted using QIAprep Spin Miniprep Kit (Qiagen). All enzymes were supplied from New England Biolabs.

### CRISPR RNP KO of tensin

The gRNA complex was prepared by mixing 1 μl of tensin 3 crRNA (100 μM; AGUC-CGCUCCCGCUCAUAG, SIGMA) and 1 μl of trRNA (100 μM; 1072533, IDT) with 98 μl of nuclease-free water to prepare a complex of 100 μM final concentration. The complex was heated for 5 minutes at 95°C to assemble the complex and was then left to cool on the bench to room temperature. The transfection complex was prepared by mixing gRNA complex with 1 μΜ Cas9 nuclease V3 (1081058; IDT) at a ratio of 1,3:1 in Opti-MEM and incubate for 5 minutes at room temperature. The gRNA:Cas9 complex was mixed with lipofectamine 2000, used per the manufacturer instructions and incubated for 20 minutes at room temperature. Reverse transfection was performed by adding the transfection complex to freshly plated cells (RNP final concentration, 10 nM). Limited dilution was performed after 48 hours of incubation with the transfection complex to collect individual cells which were expanded and tested for tensin 3 expression by Immunofluorescence and western blot. To assess CRISPR efficiency, DNA was extracted from the cells and sequenced. The sequences were uploaded on ICE to assess how many cells were missing the targeted sequence.

### Generation of stable cell lines

U2OS cells were co-transfected with pCDH-TagBFP-T2A-myc-BirA* or pCDH-TagBFP-T2A-myc-BirA*-tensin3 along with a puromycin vector at a 5:1 ratio. Twenty-four hours after transfection, cells were trypsinized and diluted at a ratio 1:5 before replated. Stable cells were selected with 1.5 μg ml^−1^ puromycin (1 mg ml^−1^ stock; Invitrogen) 24 hours after replating. Upon colony formation, individual blue clones were isolated using colony rings (Sigma) and screened by cell imaging and western blot, before expanded. Stable cells were then supplemented with 0.5 μg ml^−1^ puromycin until cell line was fully established and expanded to 10cm dishes. Established stable cells were habituated in either heavy (H) or control light (L) isotopic labelled DMEM for SILAC (ThermoFisher) for 14 days (6 passages) supplemented with 10% dialyzed FBS (ThermoFisher), 1% Pen/Strep, 143 mg ml^−1^ Lysine and 83 mg ml^−1^ Arginine (K8R10 ‘Heavy’ or K0R0 ‘Light’). On the last passage step, 2-4 dishes were kept for each of the control and Tensin3 cell lines, depending on how strong the expression was. In our case 4 10cm-dishes of BFP-myc-BirA*-tensin3 were used and 2 10cm-dishes of BFP-myc-BirA* (due to stronger expression). We performed the experiment in both directions with BFP-myc-BirA*-tensin3 adapted in “heavy” labelled medium combined with BFP-myc-BirA* adapted in “light” labelled medium and vice versa.

### BioID and NeutrAvidin affinity capture

Labelled cells with heavy or light SILAC medium were in-situ labelled with D-biotin (100 μM; invitrogen, B20656) overnight (16 hours). The next day the cells were washed three times with phosphate-buffered saline (PBS) (5 minutes, room temperature) to remove the excess biotin. Cell were lysed in 500 μl of lysis buffer (25 mM Tris (pH 8.0), 100 mM KCl, 5% glycerol, 0.5% Triton X−100, 0.5% deoxycholate (DOC), 1 mM EDTA, and protease inhibitor cocktail; Roche) per 10 cm dish. The cells were disrupted using a p1000 tip. The lysates were then centrifuged at maximum speed (21,130g) for 1 minute to remove nuclei, and the supernatant was transferred to a fresh 1,5 ml tube. The lysates were made up to 0.2% SDS and sonicated for 20 seconds at 15% power, using a tapered tip before they were centrifuged (13,200 rpm) for 10 minutes at 4°C. At this stage, the heavy-labelled samples were mixed with light-labelled samples and added to 100 μl Neutravidin beads slurry, which have been previously equilibrated with lysis buffer, and incubated with rolling overnight at 4°C. The next day the samples were centrifuged at 4,000 rpm for 2 minutes to collect the beads. Supernatant was removed and the beads were washed with 500 μl lysis buffer + 0.2% SDS, for 15 minutes at 4°C. The beads were then washed twice in wash buffer (25 mM Tris (pH 8.0), 50 mM NaCl, 1% SDS, and protease inhibitor cocktail) for 10 minutes at room temperature and 4°C respectively. Proteins bound to Neutravidin beads were eluted in 80 μl of 1× lithium dodecyl sulfate (LDS) sample buffer (Novex NuPAGE) with 1 mM biotin, by heating at 95°C for 5 minutes. This was removed and replaced by 40 μl of distilled water which was heated with the beads at 95°C for another 5 minutes to rinse the beads. The 40 μl were then added to previous 80 μl of elute and they were reduced to 40 μl under vacuum to concentrate the sample. Purified proteins (35μl) were run on a 10% SDS-PAGE gel (1mm) and stained with colloidal Coomassie Brilliant Blue (CBB, Novex, Invitrogen). Those with a weight >28 kD (Streptavidin dimer) were excised processed for trypsin digestion using in-gel digestion procedures (Shevchenko et al. 2006) and interaction candidates were identified using mass spectrometry. Protein enrichment was calculated as a Heavy/Light peptide ratio and used for analysis as described in (Dong et al. 2016).

### Protein identification by SILAC mass spectrometry

Tryptic peptides were analysed using an EASY-nLC 1000 coupled to a Q ExactiveTM Hybrid Quadrupole-Orbitrap (Thermo Fisher Scientific). The peptides were resolved and separated on a 50cm analytical EASY-Spray column equipped with pre-column over a 120 min gradient ranging from 8 to 38% of 0.1% formic acid in 95% acetonitrile/water at flowrate of 2 00nl/min. Survey f ull scan MS spectra (m/z 310–2000) were acquired with a resolution of 70k, an AGC target of 3×106 and a maximum injection time of 10ms. Top twenty most intense peptide ions in each survey scan were sequentially isolated to an ACG target value of 5e4 with resolution of 17,500 and fragmented using normalized collision energy of 25. A dynamic exclusion of 10s and isolation width of 2 m/z were applied. Data analysis was performed with MaxQuant (Cox and Mann 2008) version 1.6.0.1 using default settings. Database searches of MS data used Uniprot human fasta (2020 Jan release, 96817 proteins) with tryptic specificity a llowing m aximum t wo m issed cleavages, two labelled amino acids, and an initial mass tolerance of 4.5 ppm for precursor ions and 0.5 Da for fragment ions. Cysteine carbamidomethylation was searched as a fixed modification, and N-acetylation and oxidized methionine were searched as variable modifications. Maximum false discovery rates were set to 0.01 for both protein and peptide. Proteins were considered identified when supported by at least one unique peptide with a minimum length of seven amino acids. Proteins with a ratio count <4 were excluded from the filtered data (Supp. Table 1), while mitochondrial proteins, histones and biotin-related proteins were excluded from the potential binding partner list (Fig. 2b). The mass spectrometry proteomics data have been deposited to the ProteomeXchange Consortium via the PRIDE (Perez-Riverol et al. 2019) partner repository with the dataset identifier PXD026343.

### Microscopy

#### Live-cell imaging

Images of transfected vinKO MEFs (prepared as described above) were acquired on a spinning disk confocal microscope (CSU-X1, Yokagowa) supplied by Intelligent Imaging Innovations, Inc. (3i) equipped with a motorized XYZ stage (ASI) maintained at 37°C, using a 100x/1.45 Plan Apo oil objective (Zeiss) and an Evolve EMCCD camera (Photometrics). One hour prior to imaging the medium was changed to pre-warmed Ham’s F–12 medium supplemented with 25 mM HEPES buffer, 1% FCS, 1% penicillin/streptomycin and 1% L-glutamine, with 5 mM Mn^2+^ added as appropriate. The 488nm and 561nm lasers were controlled using an AOTF through the laserstack (Intelligent Imaging Innovations (3I)).

#### Fluorescence recovery after photobleaching (FRAP)

Transfected NIH3T3 fibroblasts were incubated overnight at 37°C. The cells were placed in the microscope chamber at 37°C for 1 hour prior to imaging, to ensure they were in equilibrium.Images were collected on a Leica Infinity TIRF microscope using a 100x/1.47 HC PL Apo Corr TIRF Oil objective with a 488nm diode TIRF laser (with 100% laser power and a penetration depth of 120nm), an ORCA Flash V4 CMOS camera (Hamamatsu) with 400ms exposure time and camera gain of 2, and a Leica QwF-S-T filter c ube. Leica LAS X software was used t o bleach 5-7 adhesions per cell (using regions of interest manually drawn around adhesions); 3 pre-bleach images were acquired, followed by one image every 10 seconds for 5 minutes post-bleach. Movies were analysed using FIJI-ImageJ.

#### Fixed-sample imaging

Images of fixed s amples in PBS were acquired at room temperature using a Zeiss AxioObserver Z1 wide-field microscope equipped with a 100×/1.4-NA oil objective and an Axiocam MRm camera, controlled by Zeiss Axiovision software. Samples were illuminated using a mercury bulb; specific band-pass filter sets were used to prevent bleed through from one channel to the next (for GFP, 38HE (Zeiss); for mCherry, 43HE (Zeiss)). Images of fixed U2OS samples in PBS were acquired at room temperature using a Leica DM6000 B wide-field microscope equipped with a 63x/1.4 oil objective and a photometrics coolsnap EZ camera, controlled by metamorph software. Samples were illuminated using the external light source for fluorescence excitation; Leica EL6000 (mercury metal halide lamp); specific band-pass filter sets were used to prevent bleed through from one channel to the next (for GFP, L5 (Leica); for mCherry, TX2 (Leica); for DAPI, A4 (Leica)).

### Analysis of cell-matrix adhesions

Cell-matrix adhesion size, number and percent area were quantified u sing FIJI-ImageJ (Schindelin et al. 2012) as described previously (Atherton et al. 2015), by subtracting background signal using a rolling ball algorithm, followed by thresholding to select particles. Similar analysis was performed to quantify fibronectin fibres. The distance of adhesions or fibronectin fibrils from the cell edge was quantified by using a region of interest drawn around the cell periphery, which was used to create a Euclidean distance map (EDM) using the Distance Map function in imageJ. Adhesions or fibronectin fibrils were thresholded as above and used to create masks that were applied to the EDM. The mean pixel intensity in each adhesion/fibril mask applied to the EDM gives the distance from the cell edge.

### *In vitro* peptide binding assays

#### Peptide and Protein Preparation

Recombinant wild-type mouse talin1 fragment R11 (residues 1,974–2,140) was previously cloned into pET151/D-TOPO expression vector (Gingras et al. 2009). Protein was produced in BL21 STAR (DE3) cultured in Luria-Bertani or 2×M9 minimal medium containing 1 g/l 15N-labeled NH4Cl, and purified using nickel-affinity chromatography followed by ion exchange. Synthetic peptide corresponding to TNS3 H1 fragment (residues 692-712) was purchased from ChinaPeptides (Shanghai). The peptide was dissolved in milliQ water at concentration of 10 mM and pH adjusted to nerutral by adding NaOH. Required amout of the stock peptide solution was added to buffer for ITC or NMR sample of talin for the titration experiments.

#### NMR Spectroscopy

NMR spectra were collected on Bruker Neo 800 MHz spectrometers equipped with TCI CryoProbe. Experiments were performed at 298 K in 20 mM sodium phosphate (pH 6.5) and 50 mM NaCl with 5% (v/v) 2H2O. Spectra were processed with TopSpin (Bruker).

#### Isothermal titration calorimetry (ITC)

ITC experiments were performed using an ITC-200 (Microcal). ITC titrations were performed in 20 mM sodium phosphate (pH 6.5) and 50 mM NaCl and 0.5mM TCEP (tris-carboxyethyl-phosphine) at 25oC. Data were integrated and fitted to a single-site-binding equation using Origin 7 software with integrated ITC module (Microcal).

### Graphs and statistical analysis

All graphs were made using Prism 8 (GraphPad). Statistical analyses were performed using Prism 8 (GraphPad). Where appropriate, statistical significance between two individual groups was tested using a (two-tailed) t-test. To test for significance between two or more groups, a One-way ANOVA was used with a Holm-Sidak’s multiple comparison test with a single pooled variance. Data distribution was tested for normality using a D’Agostino & Pearson omnibus K2 test; a p-value >0.05 was used to determine normality.

---

**Supplemental Fig. 1.**
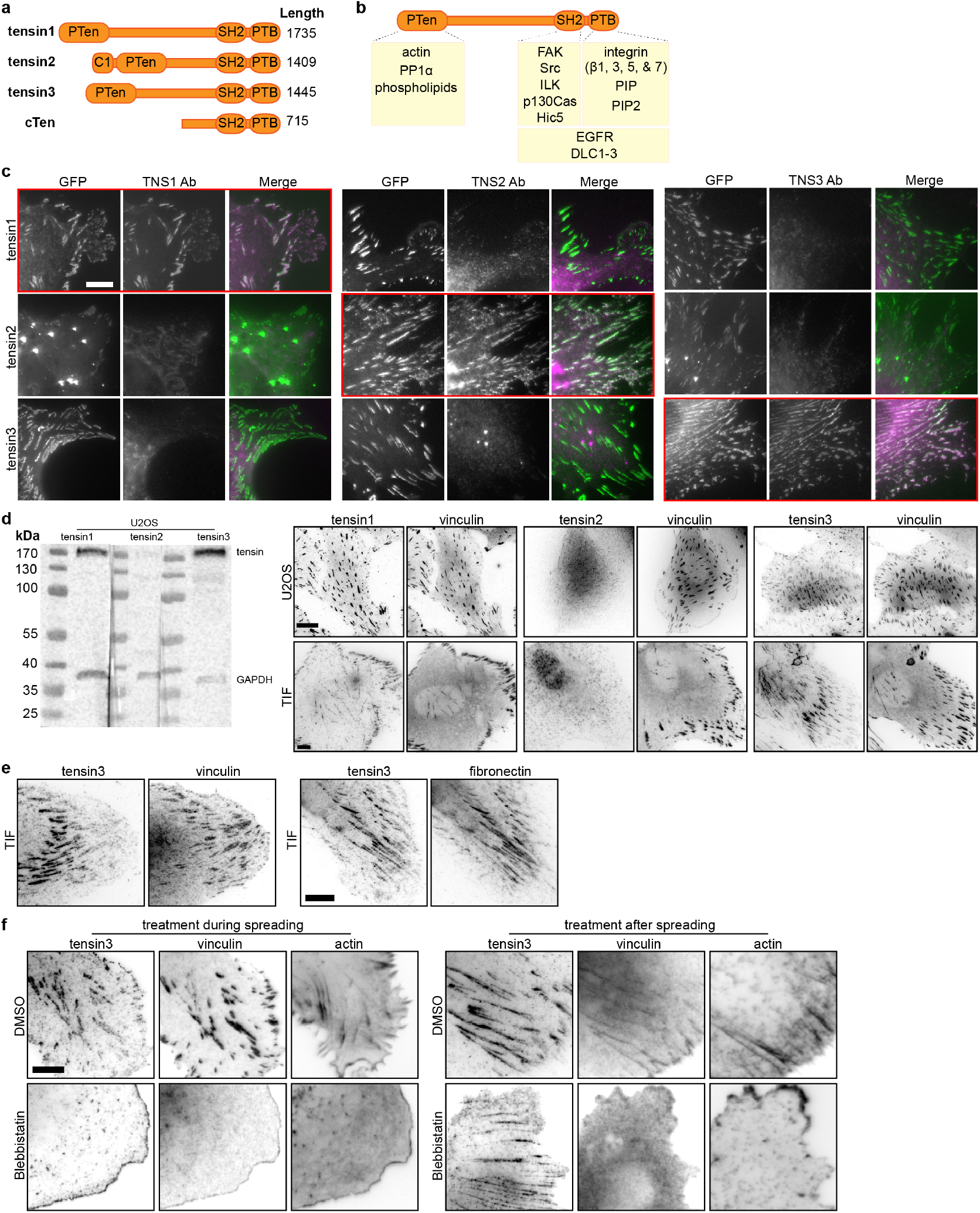
Characterisation of tensin specific antibodies and tensin localisation. **a**, Schematic of the four human tensin family members. Tensin1-3 have an N-terminal region homologous to phosphatase and tensin homolog (PTEN). Tensin2 has an N-terminal lipid-binding C1 domain. All four family members have C-terminal Src-homology 2 (SH2) and phosphotyrosine-binding (PTB) domains. The middle regions are largely unstructured with little amino acid sequence homology between family members. **b**, Schematic showing various reported interaction partners (reviewed in (Liao and Lo 2021)) of the tensin family, including: cytoplasmic domains of beta integrin subunits (β1, β3, β5, β7) with the PTB domain that reportedly supports integrin activation (Calderwood et al. 2003; Katz et al. 2007; McCleverty, Lin, and Liddington 2007); the SH2 domain with p130CAS (Zhao et al. 2016); actin to the N-terminus (Lo et al. 1994); the Rho GAPs DLC1-3 to both the Nand C-termini (Shih et al. 2015; Cao et al. 2012; Liao et al. 2007; Kawai et al. 2009; Qian et al. 2007); growth factor receptors (EGFR (Cui, Liao, and Lo 2004; Katz et al. 2007), cMET (Muharram et al. 2014)) and additional FA proteins (e.g. FAK and p130Cas (Qian et al. 2007; Hall et al. 2010; Cui, Liao, and Lo 2004), Hic5 (Goreczny, Forsythe, and Turner 2018), ILK (Qian et al. 2007)) to the C-terminus. **c**, NIH3T3 mouse fibroblasts were transfected with the indicated GFP-tagged human tensin constructs and immunostained with antibodies against (human) TNS1, TNS2, or TNS3. Note that the respective antibodies detect only the expressed tensin member they are directed against (i.e. tensin1 antibody detects only over-expressed GFP-tensin1, tensin2 antibody only the expressed GFP-tensin2 and the tensin3 antibody only the GFP-tensin3); indicated by red boxes). **d**, Expression of tensin1, 2 and 3 in U2OS and TIF cells evaluated by western blot and immunofluorescence. **e**, Left panel: TIF cell stained for vinculin and tensin3, note that tensin3 localises more to the centrally located adhesions and vinculin to the peripheral adhesions; right panel: TIF cell stained for tensin3 and fibronectin, note the similar localisation. **f**, Left panels: TIF cells were treated in suspension with blebbistatin (50 μM) or an equivalent volume of DMSO, for 60 minutes. Cells were fixed after spreading on fibronectin-coated glass for 60 minutes. Note the absence of tensin3- or vinculin-positive structures in blebbistatin-treated cells. Right panels: TIF cells cultured overnight on fibronectin-coated glass were treated with blebbistatin (50 μM) or an equivalent volume of DMSO, for 45 minutes prior to fixation. Scale bars in c-f indicate 10 μm.

**Supplemental Fig. 2.**
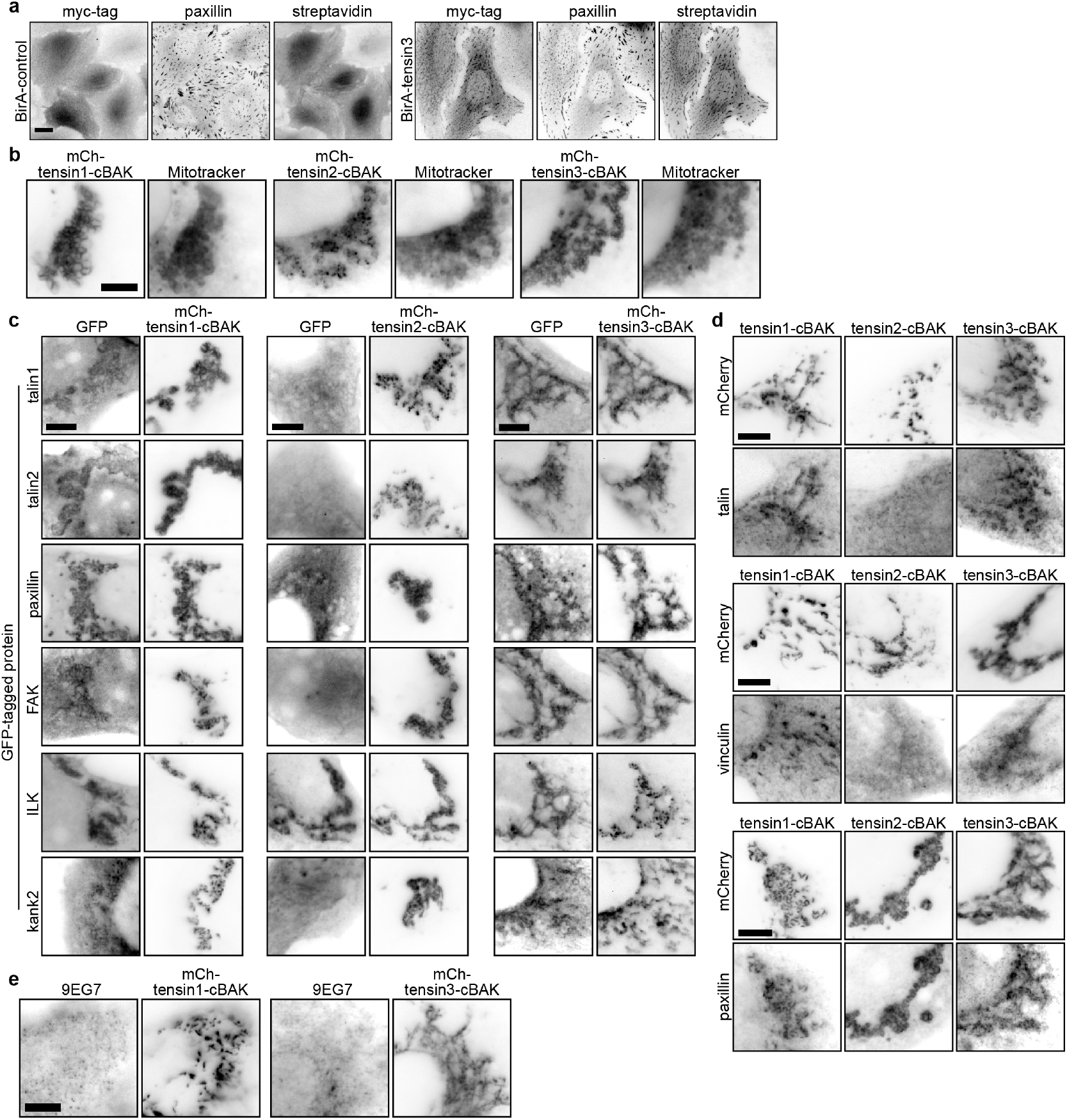
The mitochondrial targeting system confirms tensin interactions with other FA proteins. **a**, U2OS cells stably expressing BirA-control or BirA-tensin3 were incubated with biotin for 16 hours before being fixed and stained for myc and paxillin and biotinylated proteins (using fluorophore (Dylight 488)-conjugated streptavidin). Scale bar indicates 10 μm. **b**, The C-terminus of tensin1, 2 or 3 was fused to the short mitochondrial targeting sequence from the outer mitochondrial membrane protein BAK (cBAK), with an N-terminal mCherry (mCh) tag. When expressed in NIH3T3 cells each of these constructs co-localise with the mitochondria-specific dye MitoTracker. **c**. Representative images of NIH3T3 cells co-expressing the indicated GFP-tagged cell-matrix adhesion protein together with the indicated mCh-tensin-cBAK construct. **d**, Representative images of NIH3T3 cells expressing mCh-tensin-cBAK constructs immunostained for endogenous talin, vinculin or paxillin. **e**, Representative images of NIH3T3 cells expressing mCh-tensin1-cBAK or mCh-tensin3-cBAK immunostained for active β1 integrin (9EG7 stain) reveals the absence of active integrins. Scale bars in b-e indicate 5 μm.

**Supplemental Fig. 3.**
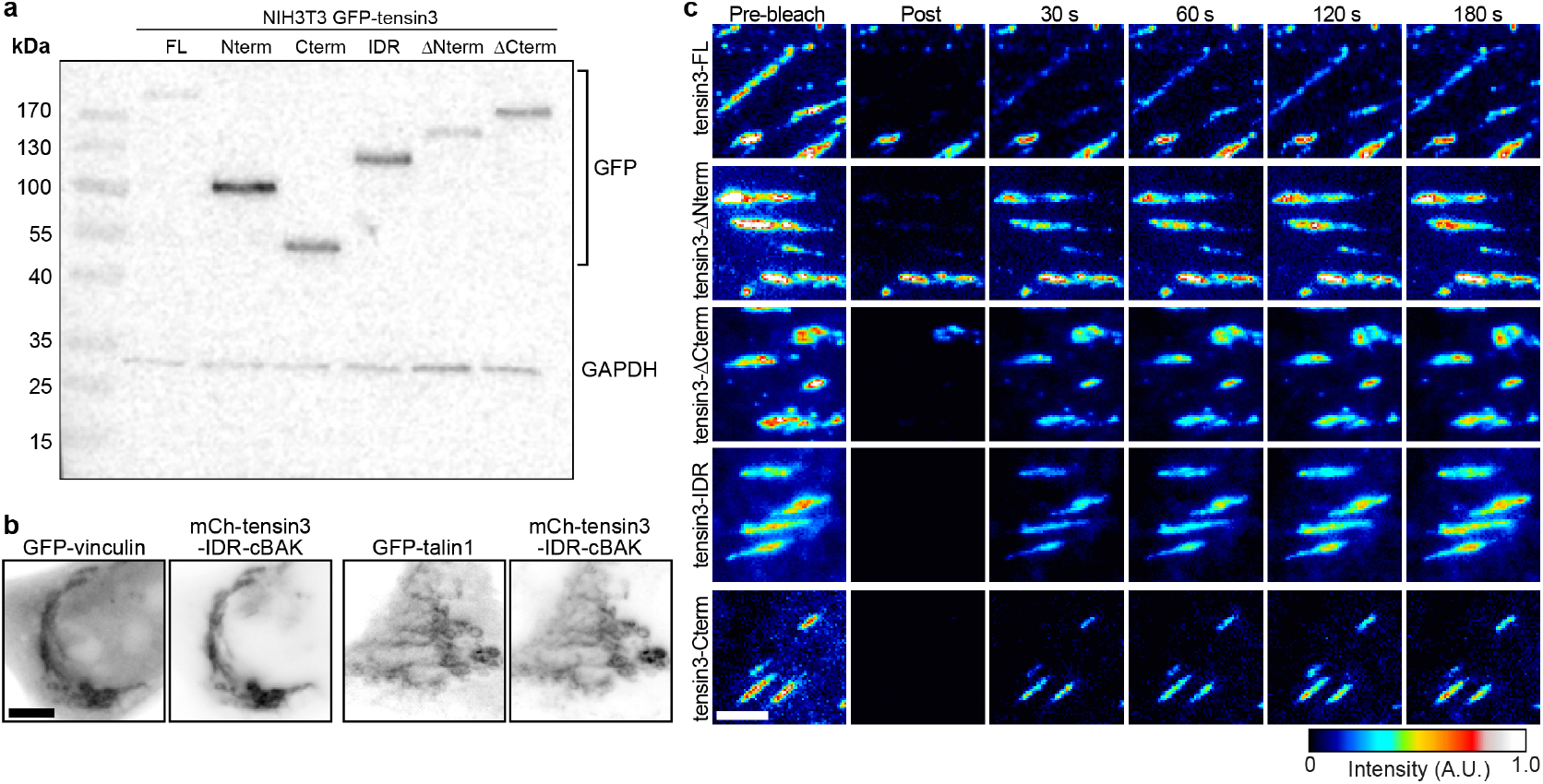
**a**, Western blot of tensin3 constructs N-terminally tagged with GFP expressed in NIH3T3 cells. **b**, Co-expression of mCh-tensin3-IDR-cBAK with either GFP-vinculin or GFP-talin1 in NIH3T3 cells reveals either protein is recruited to the tensin3 IDR at mitochondria. Scale bar indicates 5 μm. **c**, Representative images from FRAP experiments in NIH3T3 cells. Scale bar indicates 3 μm.

**Supplemental Fig. 4.**
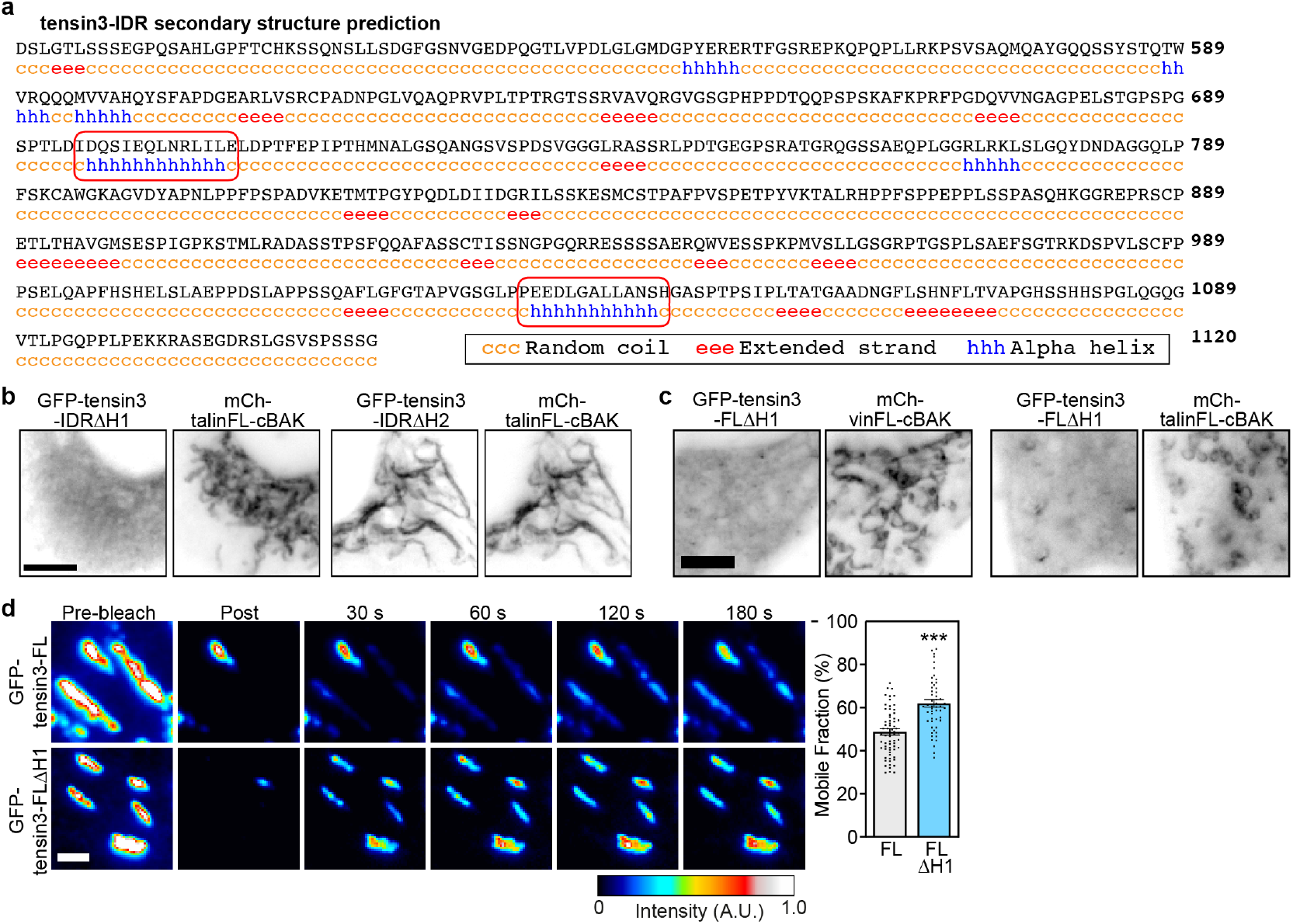
**a**, Secondary structure prediction of the tensin3 IDR (aa490–1120). Red boxes indicate the H1 and H2 regions. **b**, mCh-talinFL-cBAK was expressed in NIH3T3 cells together with a tensin3-IDR construct lacking either the H1 (GFP-tensin3-IDRΔH1) or H2 (GFP-tensin3-IDRΔH2) motif. Note the absence of co-localisation between GFP-tensin3-IDRΔH1 and mCh-talinFL-cBAK, indicating that the H1 motif is responsible for this interaction. **c**, A tensin3 construct lacking the H1 motif (GFP-tensin3-FLΔH1) co-expressed in NIH3T3 cells with either mCh-vinFL-cBAK or mCh-talinFL-cBAK is not recruited to mitochondria under either condition. Scale bars in b, c, indicate 10 μm. **d**, Representative images from FRAP experiments performed in NIH3T3 cells expressing either GFP-tensin3-FL or GFP-tensin3-FLΔH1; note the faster recovery of GFP-tensin3-FLΔH1 compared to GFP-tensin3-FL, and the higher mobile fraction. Error bars are S.E.M, n = 63 (GFP-tensin3-FL) or 51 (GFP-tensin3-FLΔH1) adhesions from 7 cells; *** indicates p<0.001 (t-test); scale bar indicates 2 μm.

**Supplemental Fig. 5.**
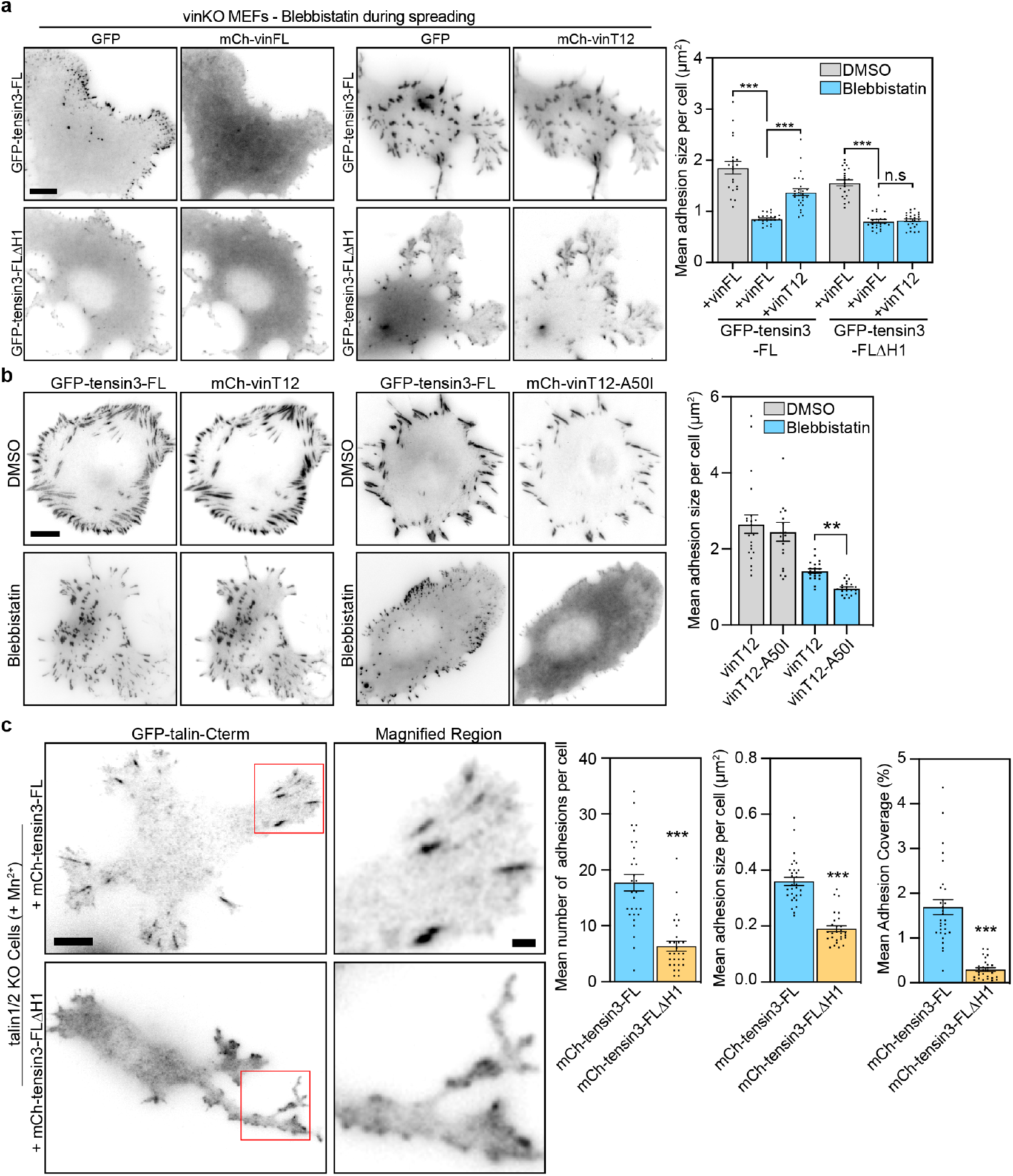
**a**, Representative images of vinculin-null cells (vinKO MEFs) co-expressing GFP-tensin3-FL or GFP-tensin3-FLΔH1 together with either mCh-vinFL or mCh-vinT12, treated in suspension with blebbistatin (50 μM) or an equivalent volume of DMSO, for 60 minutes. Cells were fixed after spreading on fibronectin-coated glass for 60 minutes. Quantification of mean adhesion size per cell using the GFP signal shows that co-expression of active vinculin (mCh-vinT12) can rescue formation of GFP-tensin3 positive adhesions under tension-free (blebbistatin-treated) conditions; this phenotype is not seen in cells expressing GFP-tensin3-FLΔH1. Error bars are S.E.M; n = 19-24 cells, *** indicates p<0.001 (ANOVA), n.s. indicates not significant. **b**, Representative images of vinculin-null cells (vinKO MEFs) co-expressing GFP-tensin3-FL together with either mCh-vinT12 or mCh-vinT12-A50I, treated in suspension with blebbistatin (50 μM) or an equivalent volume of DMSO, for 60 minutes. Cells were fixed after spreading on fibronectin-coated glass for 60 minutes. Quantification of mean adhesion size per cell using the GFP signal shows that inhibiting the vinculin-talin interaction (vinT12-A50I expression) blocks the formation of large GFP-tensin3-FL positive adhesions in tension-free conditions. Error bars are S.E.M; n = 19-21 cells, ** indicates p<0.001 (Kruskal-Wallis test). Scale bars in a, b indicate 10 μm. **c**, Representative images of talin1/2 KO cells co-expressing GFP-talin-Cterm with either mCh-tensin3-FL or mCh-tensin3-FLΔH1, fixed after 1 hour of spreading on fibronectin-coated glass in the presence of manganese (5 mM). Red box indicates magnified region; scale bar indicates 5 μm; scale bar in magnified region indicates 2 μm. Quantification of adhesions using the GFP signal shows that adhesion structures are larger and more numerous in cells expressing mCh-tensin3-FL compared to those expressing mCh-tensin3-FLΔH1. Error bars are S.E.M; n = 29 (mCh-tensin3-FL) or 28 (mCh-tensin3-FLΔH1) cells; *** indicates p<0.001 (t-test).

**Supplemental Fig. 6.**
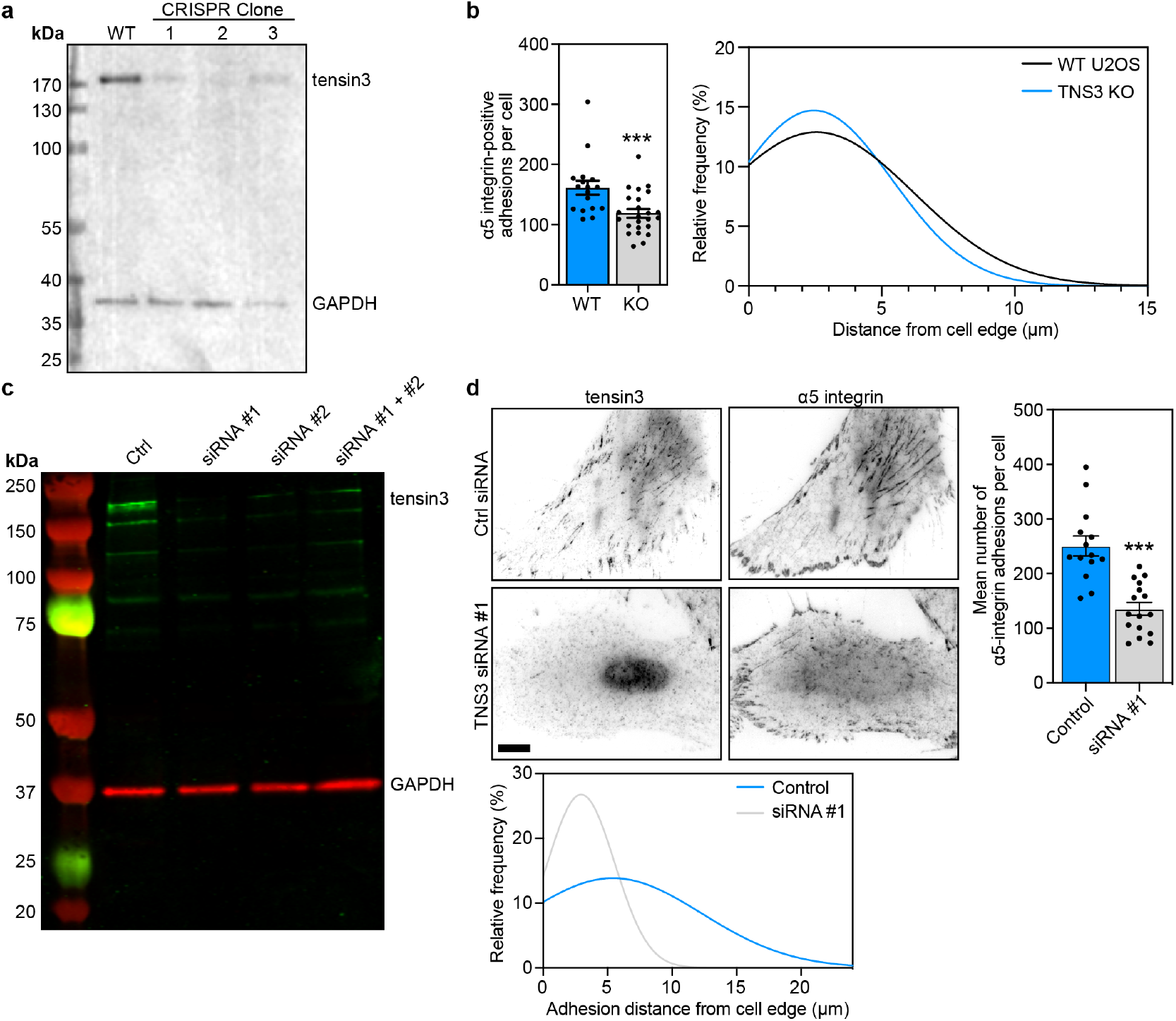
Depletion of tensin3 using CRISPR and siRNA. **a**, Generation of TNS3 KO cells using CRISPR; western blot shows endogenous tensin3 levels in WT and 3 clonal populations generated following CRISPR. Clone #2 was a complete KO line (all cells were depleted of TNS3); clones #1 and #3 were a mixed population of KO and WT cells. **b**, WT or TNS3 KO (clone #3) U2OS cells were spread overnight on fibronectin-coated glass prior to fixation and immunostaining for α5 integrin. TNS KO cells (clone #3) have reduced α5 integrin-positive adhesions; error bars are S.E.M, n = 17 (WT) and 24 (KO) cells; *** indicates p<0.001 (Mann-Whitney test). Gaussian distribution of the distance of all detected α5 integrin-positive structures from the cell periphery. Data are pooled from three independent repeats. **c**, Western blot of endogenous tensin3 in TIFs after 2 rounds of transfection with either scrambled siRNA (Ctrl) or two different oligos targeting tensin3 either separately (siRNA #1 or siRNA #2) or pooled together (siRNA #1 + #2). Strongest knockdown was achieved with siRNA #1 alone. **d**, Representative images of TIFs after siRNA-mediated knockdown of tensin3 as described above. Cells were cultured on fibronectin-coated glass overnight prior to fixation, then immunostained for α5 integrin. Quantification shows that tensin3 knockdown reduced the number of α5 integrin-positive adhesions. Errors bars are S.E.M; n = 14 (Control) or 16 (siRNA #1) cells; *** indicates p<0.001 (t-test). Gaussian distribution of the distance of the α5 integrin-positive structures from the cell periphery shows that TNS3 knockdown in TIFs reduced the number of centrally-located α5 integrin-positive adhesions.

**Supplemental Fig. 7.**
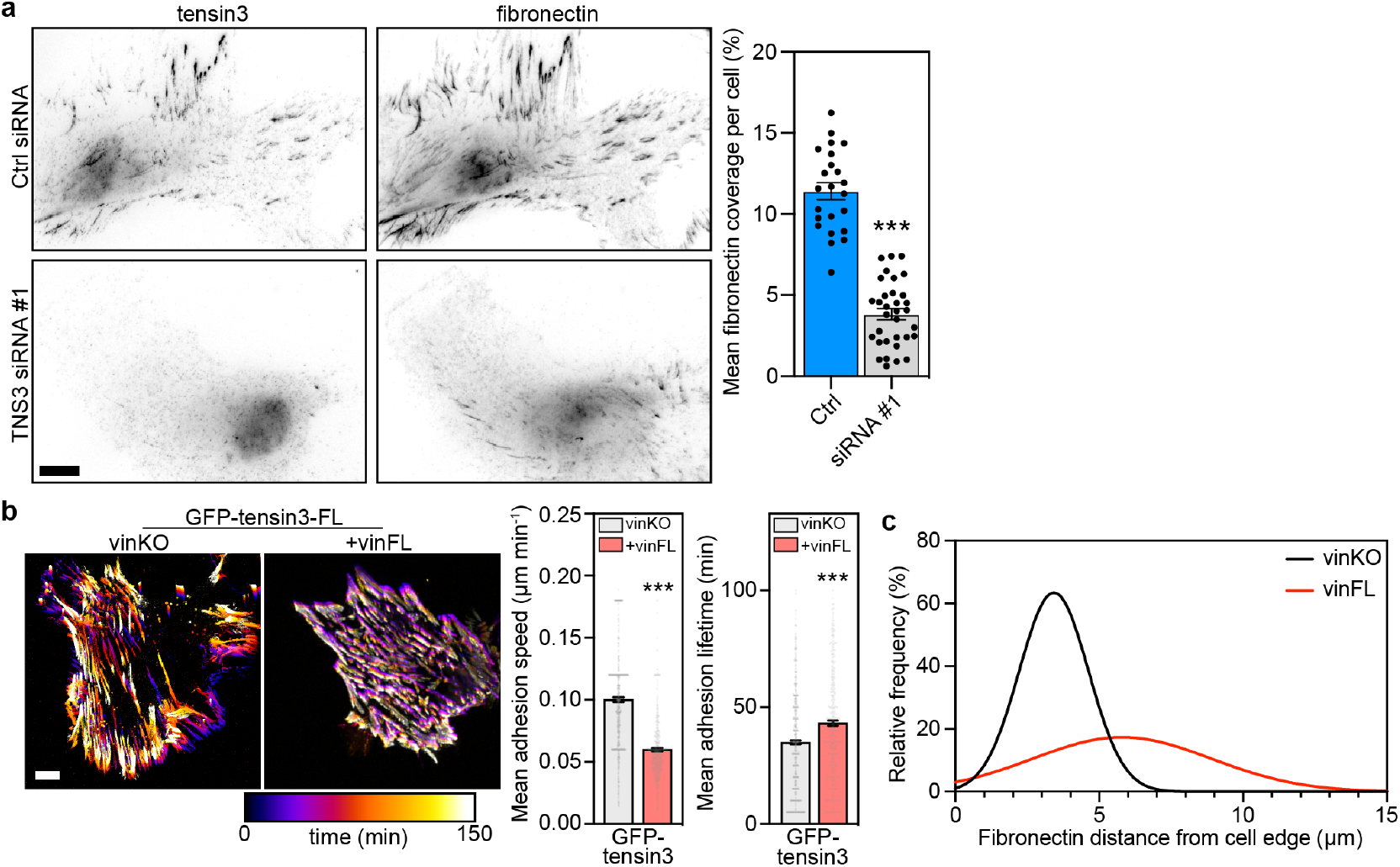
**a**, Representative images of TIFs after siRNA-mediated knockdown of tensin3. Cells were cultured on fibronectin-coated glass overnight prior to fixation, then immunostained for fibronectin. Quantification shows that tensin3 knockdown reduced the number of fibronectin fibrils. Errors bars are S.E.M; n = 23 (Control) or 33 (siRNA #1) cells; *** indicates p<0.001 (t-test). **b**, Live-cell imaging of vinKO MEFs expressing GFP-tensin3-FL with or without mCh-vinFL co-expression; temporal colour maps of adhesion movement obtained from the GFP signal show that tensin3-positive adhesions in cells without vinculin are more dynamic. Images were acquired every 5 min for 2 hr. Quantification of mean adhesion speed and lifetime from time-lapse movies of vinKO MEFs expressing GFP-tensin3-FL with or without mCh-vinFL co-expression. Error bars are S.E.M, n numbers are pooled from 6 (vinKO) or 10 (+vinFL) cells; *** indicates p<0.001 (t-test). **c**, Gaussian distribution of the mean distance of fibronectin (FN) fibres from the cell periphery. Cells without vinculin have fewer centrally-located FN fibrils compared to cells expressing mCh-vinFL. Scale bars in a-c indicate 10 μm.

## Notes

### Competing Interest Statement

The authors have declared no competing interest.

### Summary of Updates

Description of figure 7c in the Results was previously absent

